# Protein identification using cryo-EM and artificial intelligence guides improved sample purification

**DOI:** 10.1101/2024.09.11.612515

**Authors:** Kenneth D. Carr, Dane Evan D. Zambrano, Connor Weidle, Alex Goodson, Helen E. Eisenach, Harley Pyles, Alexis Courbet, Neil P. King, Andrew J. Borst

## Abstract

Protein purification is essential in protein biochemistry, structural biology, and protein design. It enables the determination of protein structures, the study of biological mechanisms, and the biochemical and biophysical characterization of both natural and de novo designed proteins. Despite the broad application of various protein purification protocols, standard strategies can still encounter challenges, such as the unintended co-purification of unknown contaminants alongside the target protein. In particular, co-purification issues pose significant challenges for designed self-assembling protein nanomaterials, as it is difficult to determine whether unexpected observed geometries represent novel assembly states of the designed system, cross-contamination from other assemblies, or native proteins originating from the expression host. In this study, we assessed the ability of an automated structure-to-sequence pipeline to unambiguously identify an unknown co-purifying protein found across several purified designed protein samples. Using cryo-electron microscopy (Cryo-EM), ModelAngelo’s sequence-agnostic automated model-building feature, and the Basic Local Alignment Search Tool (BLAST), we identified the unknown protein as dihydrolipoamide succinyltransferase (DLST). This identification was further confirmed by comparing the cryo-EM data with available DLST structures in the Protein Data Bank (PDB) and AlphaFold 3 predictions from the top BLAST hits. The clear identification of DLST informed our subsequent literature search and led to the rational modification of our protein purification protocol, ultimately enabling the exclusion of the contaminant from preparations of our target nanoparticle. This study demonstrates the successful application of a structure-to-sequence workflow, integrating Cryo-EM, ModelAngelo, protein BLAST, PDB structures, and AlphaFold 3 predictions, to identify and remove an unknown protein from a purified sample. It also highlights the broader potential of integrating Cryo-EM with AI-driven tools for accurate protein identification across various samples and contexts within protein science.

**Highlights:**

- An unknown protein was consistently found in multiple de novo designed protein samples.
- The protein was identified as dihydrolipoamide succinyltransferase (DLST) using Cryo-EM, ModelAngelo and BLAST, and further verified using AlphaFold 3 and the PDB.
- Identification enabled rational modification of the purification protocol to exclude the contaminant.
- This method enables accurate protein identification without requiring near-atomic resolution or prior sequence and structural data, making it broadly applicable to various areas of protein science.

**Graphical Abstract:** 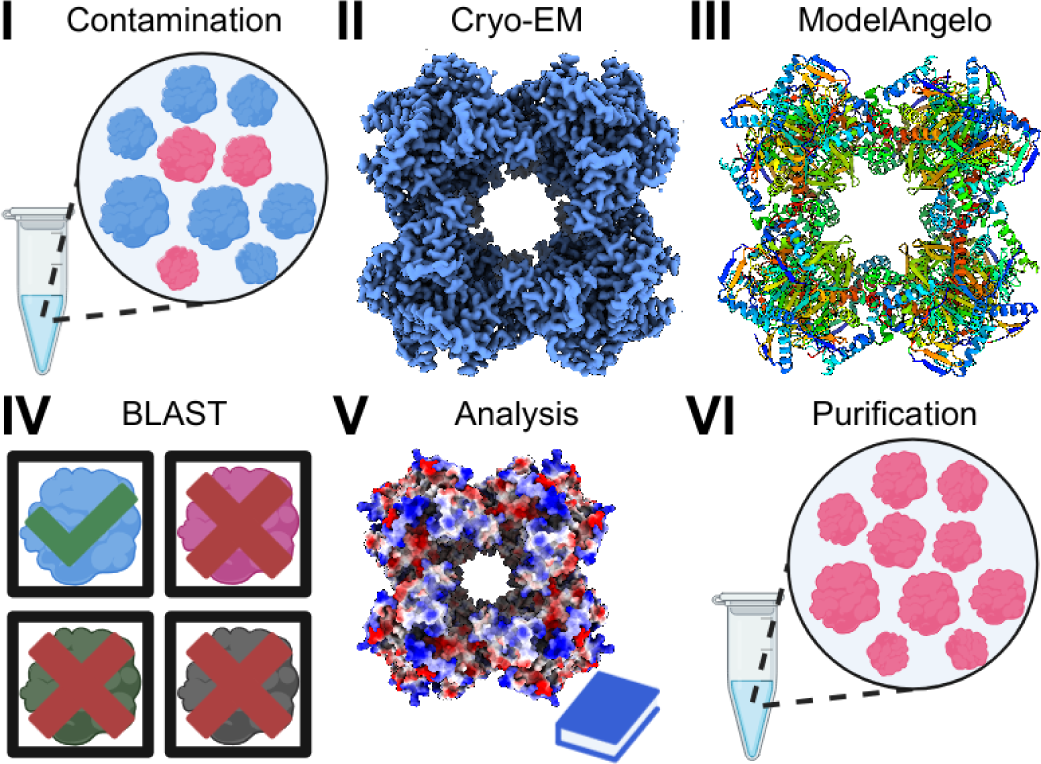

## Introduction

Understanding the details of a complex system is crucial when designing an experimental protocol, especially in the context of protein purification—a process fundamental to various scientific disciplines. The significance of protein purification lies in its ability to isolate recombinant proteins for downstream high-resolution structure determination, exploration of biochemical mechanisms, development of novel therapeutics, and the characterization of computationally designed proteins with specific functions. Despite their widespread use, protein purification protocols still face challenges, such as the co-purification of contaminant proteins from unknown origins [1]. This is particularly problematic in computational protein design, where it is often unclear whether these unknown co-purifying proteins represent off-target design states, cross-sample contamination, or naturally occurring proteins originating from the expression host.

In this study, we detail an experience encountered during the characterization of a novel computationally designed two component nanoparticle [2,3]. During initial purification attempts, we observed an unidentified protein co-eluting with our target nanoparticle, which was also found in several unrelated designed protein samples. To identify the protein, we employed both Cryo-EM single particle analysis and ModelAngelo. Single particle analysis in Cryo-EM is a technique that involves imaging thousands of individual biomolecules, then aligning and averaging their 2D projections to reconstruct a high-resolution 3D structure. ModelAngelo, a machine-learning tool, automates atomic model building and aids in protein identification by analyzing sequence fragments from Cryo-EM 3D structure information [4]. ModelAngelo’s ability to automatically build into cryo-EM density maps in a sequence-agnostic manner makes it particularly well-suited for the identification of unknown proteins in complex samples where manual methods would be time-consuming or prone to errors. By incorporating Cryo-EM and ModelAngelo into our diagnostic workflow, alongside protein BLAST, the PDB, and AlphaFold 3, we successfully identified the contaminant protein in our sample as dihydrolipoamide succinyltransferase (DLST) [4,5]. This unambiguous identification allowed us to rationally refine our purification protocol to eliminate the contaminant from future preparations.

The successful application of ModelAngelo in this context demonstrates the broad potential of integrating structural, bioinformatic, and AI-driven tools for the identification of unknown proteins in complex samples. This approach offers a powerful solution for overcoming persistent biochemical challenges, particularly in protein sample preparation, and highlights the value of such workflows when conventional sequence and structural data are unavailable.

## Results

### Characterization of an unknown co-eluting protein contaminant using electron microscopy

During the assembly and purification of a novel two-component protein nanoparticle with designed octahedral symmetry (detailed in a separate publication), we discovered that a significant portion of the purified sample consisted of a smaller, unidentified protein complex, which also exhibited octahedral symmetry as revealed by ns-EM **(Figure 1, Supplemental Table 1)**. This smaller octahedral complex was also observed in various unrelated samples from the same laboratory, including samples of de novo designed fibrous proteins, cyclic oligomers, and two distinct icosahedral nanoparticles **(Supplemental Figure 1A-D)**. In each of these other instances, the unknown protein was low in concentration relative to the designed protein, and thus could be largely ignored during downstream characterization and analysis. However, for the designed octahedral two-component nanoparticle discussed here, the presence of this contaminant comprised the vast majority of the purified protein mass **(Figure 1A-E, Supplemental Table 1)**. This high level of contamination greatly hindered our ability to accurately assemble and characterize the designed nanoparticle in the quantity and purity necessary for effective downstream biochemical, biophysical, structural, and functional applications. To improve the purification of this two-component nanoparticle, identifying the contaminating protein and its source became essential.

**Figure 1.**
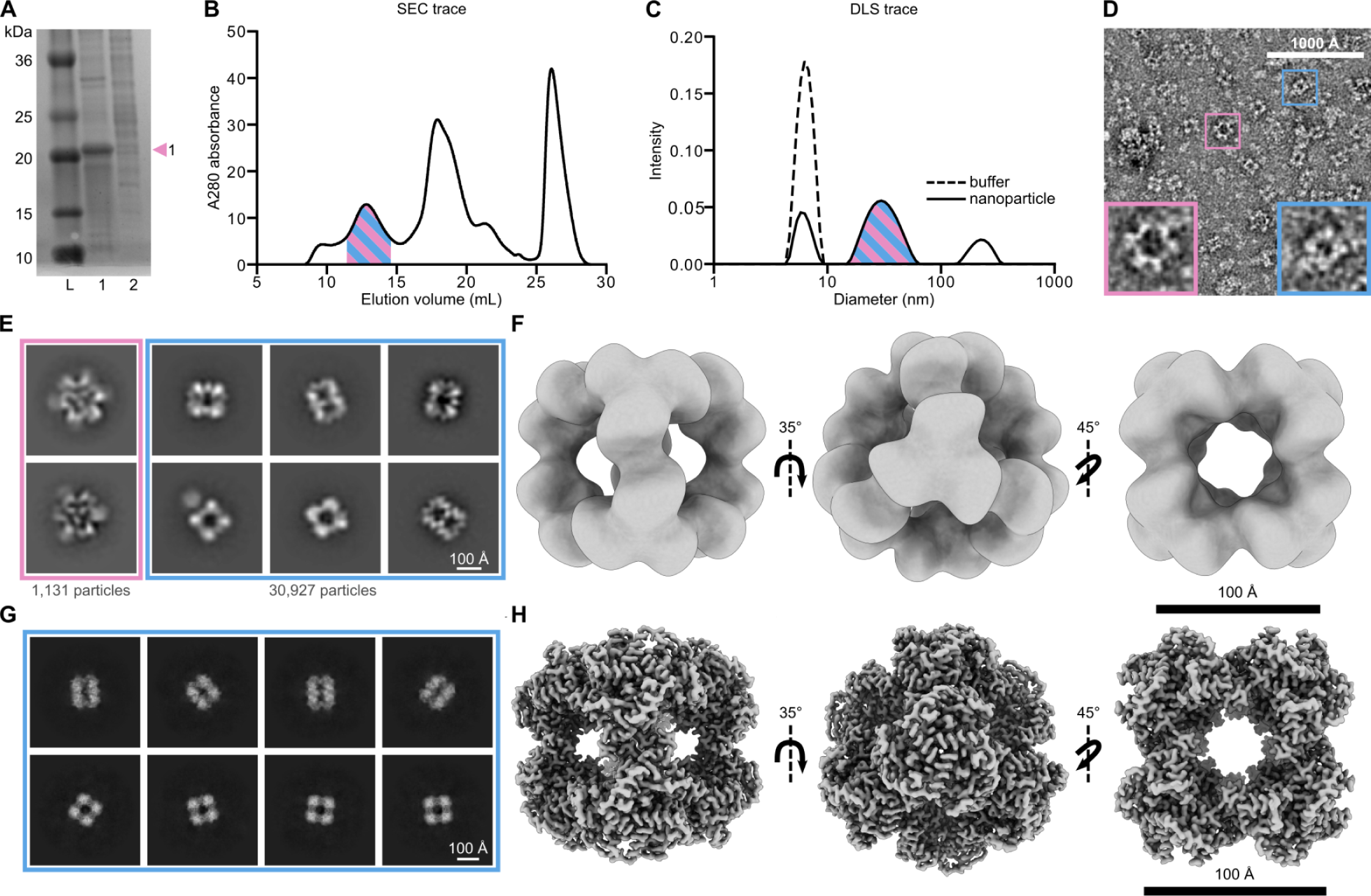
Characterization of an unknown co-eluting protein via electron microscopy. (A) SDS-PAGE of the IMAC eluate for the two protein components before mixing to form the target octahedral nanoparticle. There is a clearly defined band in lane 1 for the first protein component. The second component, shown in lane 2, is present in lower amounts and is less pure. (B) SEC trace of the assembled nanoparticle. The peak corresponding to the nanoparticle is highlighted in stripes to represent the co-elution of the contaminant with the on-target nanoparticle. (C) DLS trace of the SEC-purified sample highlighted in stripes to represent the contaminant and on-target nanoparticle diameters are not separated efficiently by DLS. The buffer (dotted line) was run as a control and contains detergent micelles found at 6.30 nm in diameter. (D) A portion of a ns-EM micrograph of the heterogeneous sample. The pink box represents the intended nanoparticle and the blue box represents the contaminant nanoparticle. (E) 2D ns-EM class averages of the on-target and contaminant nanoparticle species. A total of 1,131 on-target nanoparticles were processed and are represented here by two 2D class averages. 30,927 particles were processed for the contaminant species and are represented here by six 2D class averages. Corresponding particle numbers are reflective of all particles of each species in the dataset, not only those of the displayed classes. (F) 3D ns-EM map along the 2, 3, and 4-fold axes of symmetry of the contaminant nanoparticle using CryoSPARC v4.4. (G) 2D Cryo-EM class averages and (H) 3D Cryo-EM map along the 2, 3, and 4-fold axes of symmetry of the contaminant nanoparticle using CryoSPARC v4.5. (Blue = off-target contaminant protein; Pink = on-target two component nanoparticle).

Importantly, the intended designed two-component nanoparticle had a predicted mass of 1.20 MDa, a diameter of 29.2 nm, and required the presence of detergent for solubilization and purification. Our purification protocol first required the two protein components to be purified independently via immobilized metal-affinity chromatography (IMAC) [1,6,7]. The purified components were initially characterized via sodium dodecyl sulfate–polyacrylamide gel electrophoresis (SDS-PAGE) before being combined to form the desired two-component nanoparticle assembly. This assembly was then purified using size exclusion chromatography (SEC) [6], resulting in a final estimated total protein yield of 97 μg.

The SEC trace of the putatively purified nanoparticle contained one relevant high molecular weight (MW) peak which eluted off the column between 10 mL and 15 mL, corresponding to a particle size greater than the 660 kDa MW standard for this column **(Figure 1B)**. This peak fraction was analyzed using dynamic light scattering (DLS) to estimate the diameter of the purified nanoparticle [8]. To account for the presence of detergent micelles in the purification buffers, we performed DLS on the detergent-containing buffer used for SEC purification as a control, alongside the protein-containing peak isolated from SEC. The DLS results showed three peaks with average diameters of 6.30 nm, 30.9 nm, and 223 nm. The smallest peak was identified as containing detergent micelles, consistent with the DLS profile of the control **(Figure 1C)**. The central peak was identified as the designed two-component nanoparticle, while the largest peak corresponded to protein and micellar aggregation **(Figure 1C)**. Subsequent ns-EM analysis of the SEC-purified sample revealed the presence of two distinct nanoparticles with diameters (measured diagonally across the four-fold axes) of approximately 18 nm and 22 nm (**Figure 1D**). The larger particle exhibited the predicted diameter and morphology for the designed two-component system, with spike-like proteins extending from an underlying octahedral nanoparticle scaffold **(Figure 1E)**. However, the predominant species observed by ns-EM was a smaller, cube-like particle lacking the distinctive spiked features of our intended design. As a result, the actual yield of the target nanoparticle was estimated at approximately 3 µg from a total of 97 µg of protein purified via SEC **(Figure 1E, Supplemental Table 1)**. Furthermore, subsequent ns-EM 3D refinements produced a map of this smaller nanoparticle that revealed an octahedral assembly profile which deviated significantly from our intended design **(Figure 1F)**.

Given the prevalence of the smaller octahedral complex compared to our designed nanoparticle, and its unexpected presence across multiple unrelated samples, we turned to Cryo-EM single particle analysis to investigate whether this complex might be another designed octahedral nanoparticle, potentially resulting from cross-contamination due to the shared use of purification equipment. Following Cryo-EM data collection, initial 2D class averages of the smaller nanoparticle revealed clearly defined secondary structural elements **(Figure 1G, Supplemental Figure 2A)**. Subsequent 3D refinement yielded a final map with a global resolution of 2.51 Å, with the local resolution ranging from 2.88 Å at the periphery to 2.44 Å in the protein core **(Figure 1H, Supplemental Figure 2C, Supplemental Table 2)** [9]. However, despite the high resolution of the map, the identity of the co-eluting nanoparticle remained unclear in the absence of known sequence or structure information.

### ModelAngelo structure-to-sequence identification of the contaminant protein

To identify the unknown nanoparticle in the absence of a corresponding amino acid sequence or prior structural information, we turned to the automated model-building software ModelAngelo [4] **(Figure 2A-B)**. ModelAngelo’s default mode uses deep learning to integrate available structural data and supplied sequence information to automatically build atomic models into Cryo-EM density maps [4]. It can alternatively operate in a sequence-agnostic manner, automatically generating and building polypeptide sequence fragments into well-resolved regions of cryo-EM density maps when sequence information is unavailable. Running ModelAngelo in this manner on our 2.51 Å Cryo-EM map generated 92 all-atom sequence fragments of varying lengths, along with secondary structure predictions that were in strong agreement with the density of the unknown nanoparticle **(Figure 2A-B)**. We aligned these output chain-fragments using ClustalOmega [10,11] to generate a multiple sequence alignment that was input into WebLogo, which produced a frequency plot displaying the relative abundance of amino acids at each position **(Figure 2A, Supplemental Figure 3A)**. The consensus sequence we generated from this was then input into Protein Basic Local Alignment Search Tool (Protein BLAST) [5], which returned a list of potential matches for the unknown protein contaminant. Surprisingly, 98 of the top 100 BLAST hits were sequences of the *E. coli* Dihydrolipoamide Succinyltransferase (DLST) catalytic domain **(Figure 2A-B, Supplemental Figure 3B)**. DLST is an octahedral subunit of the E2 component of the ɑ-ketoglutarate dehydrogenase complex (KGDHC), which plays a pivotal role in the tricarboxylic acid (TCA) cycle, converting ɑ-ketoglutarate to succinyl-Coenzyme A and reducing nicotinamide adenine dinucleotide [12–14]. Notably, DLST is a well-documented contaminant protein that originates from the *E. coli* BL21 (DE3) expression system [1, 14]. As a result, we performed a pairwise sequence alignment using EMBOSS Needle [15] between the Uniprot sequence A0A140NDX4 of DLST from *E. coli* BL21 (DE3) and the ModelAngelo-derived consensus sequence, excluding the first 170 residues of DLST that were unresolved in our cryo-EM map. The alignment revealed 60.3% identity and 69.3% similarity **(Figure 2A, Supplemental Figure 3C)** [16,17], strongly indicating that the contaminant was DLST.

**Figure 2.**
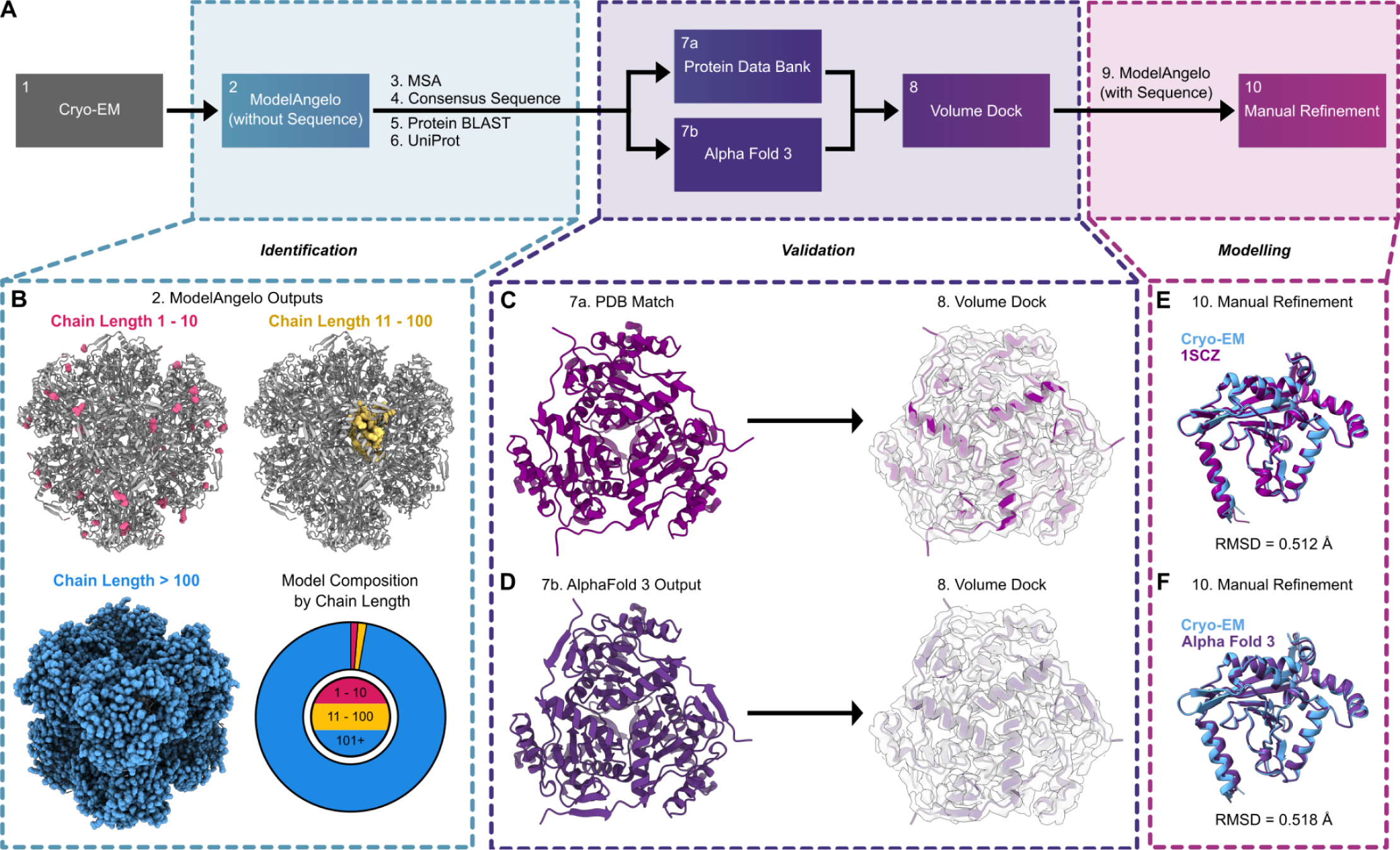
Structure-to-sequence workflow for the unambiguous identification of DLST. (A) An overview of the 10-step sequence-to-structure workflow used to identify DLST and build our atomic model used for structural analysis. (B) The sequence-agnostic ModelAngelo output using our 2.51 Å Cryo-EM map. Chains between 1 to 10 residues (pink), 11 to 100 residues (yellow), and longer than 100 residues (blue) are displayed in sphere view along the octahedral 3 fold axis of symmetry. An accompanying pie chart displays the percentage of all residues belonging to chains of those length ranges. (C-D) Published DLST crystal structure PDB:1SCZ [20] (c) and Alpha Fold 3 prediction model of the supplied DLST UniProt sequence [19] (d) each docked into the Cryo-EM density map of the unknown co-eluting protein. (E-F) Single subunit of our Cryo-EM model aligned to (E) 1SCZ (RMSD 0.512 Å) and the (F) Alpha Fold 3 model (RMSD 0.518 Å).

To further validate the identity of the unknown protein as DLST, we next compared our Cryo-EM density map with published structures of the DLST catalytic domain and AlphaFold 3-predicted structures of the DLST sequences identified by BLAST and UniProt [18,19] **(Figure 2A,C,D)**. The highest resolution crystal structure from the PDB (1SCZ; 2.20 Å) was initially selected for comparison, revealing significant agreement in alpha helix positions when docked into the cryo-EM map **(Figure 2C)** [20]. Similarly, Alpha Fold 3 (AF3) predictions using the A0A140NDX4 UniProt sequence also showed high agreement with our Cryo-EM map **(Figure 2D)** [17,19]. Together, these results unambiguously confirmed the identity of the co-eluting octahedral protein complex as DLST.

To generate our final DLST catalytic domain model, the DLST sequence derived from *E. coli* BL21(DE3) was used as the input for another round of automated model building, this time utilizing the sequence-guided feature in ModelAngelo with the UniProt sequence for DLST **(Figure 2A)** [17]. The output from ModelAngelo was then further refined to produce the final model of the DLST catalytic domain **(Figure 2A, Supplemental Figure 2B, Supplemental Figure 4)**. Aligning this final model to both the 1SCZ monomer and AF3 outputs revealed RMSD values of 0.51 Å and 0.52 Å, respectively **(Figure 2E,F)**.

### Revised purification methods for increased purity of a de novo two component protein nanoparticle

Following the identification of the contaminant protein as DLST, we turned to the literature to explore potential strategies for excluding it from our final purifications. While previous studies noted DLST’s co-purification during IMAC of many proteins, they did not offer a rationale for its co-elution, nor did any outline an effective protocol to eliminate it from the purification pipeline [1, 14]. In an effort to determine the underlying reason for DLST’s co-elution with multiple de novo protein samples, we first conducted a detailed analysis of its surface properties. Given the widespread use of *E. coli* for expressing histidine-tagged recombinant proteins, we hypothesized that the surface histidine content of DLST might contribute to its binding during IMAC purification. Clusters of surface-exposed histidine residues could potentially interact with the nickel chelating resin in a manner similar to histidine tags on recombinant proteins, leading to its unintended co-purification. [1,6,7]. However, previous reports analyzing the surface histidine content of the DLST did not find any large histidine patches to support this as a potential mechanism for co-elution from IMAC [1]. Despite the known lack of histidine clusters, we considered that non-specific interactions between single histidines on the surface of DLST may have interacted with the IMAC column **(Supplemental Figure 4A)**. To test this, we applied an elution gradient ranging from 0 M to 0.5 M imidazole during IMAC purification, aiming to isolate the individual protein components and reduce DLST’s hypothesized weak binding to the resin. However, DLST was still detected in ns-EM samples after purification **(Supplemental Figure 4B-C).** Furthermore, in samples purified with an imidazole gradient, few to no on-target two-component nanoparticle assemblies were observed **(Supplemental Figure 4C, Supplemental Table 1)**.

Given the numerous charged patches on the surface of DLST, we next tested the adjustment of sodium chloride (NaCl) concentrations in our buffers during cell lysis and IMAC purification to determine if increased ionic levels would selectively disrupt any non-specific ionic interactions occurring between DLST and the IMAC column **(Supplemental Figure 4D)** [7]. After separately purifying each protein component using IMAC with high salt buffers, followed by SEC purification of the nanoparticle, no DLST particles were detected in the sample by ns-EM **(Supplemental Figure 4E-F)**. However, while the higher NaCl concentrations eliminated DLST, it also significantly reduced the total yield of assembled on-target nanoparticles **(Supplemental Figure 4F, Supplemental Table 1).**

To exclude DLST while also preserving or improving the concentration of our on-target two-component nanoparticle, we next tested whether non-specific interactions between DLST and the on-target designed two-component nanoparticle were potentially occurring throughout purification. To do so, we opted to include excipients to all of our purification buffers—a common technique for purification of recombinant proteins to enhance yield and reduce protein aggregation [21–24]. In particular, amino acids such as glycine, threonine, arginine, glutamate, and histidine, have been successfully used as excipients to reduce non-specific protein-protein interactions, flocculation, and protein aggregation. They have also been shown to enhance protein stability in solution, particularly when working with computationally designed protein nanoparticles [25,26].

100 mM glycine and 100 mM arginine were added to all purification buffers to mitigate potential off-target interactions between our nanoparticle and DLST, which again resulted in the near-complete removal of DLST in all ns-EM micrographs of these purified samples **(Figure 3A)**. Strikingly, this approach also significantly improved the final yield of our two-component nanoparticle assembly to 220 μg of total protein **(Supplemental Table 1)**. ns-EM 2D class averages of over 70,000 particles revealed that the designed two-component nanoparticle now became the overwhelmingly dominant species, making up over 99.5% of the total particles—a 68-fold increase in on-target purified protein yield with excipients used throughout purification **(Supplemental Table 1)**. In contrast, under standard purification conditions without excipients, the target nanoparticle accounted for only 3.5% of purified particles **(Figure 3B-C, Supplemental Table 3)**. To our knowledge, this is the first reported method for effectively removing DLST as a co-purifying protein contaminant during recombinant protein production from E. coli—a challenge that, if left unaddressed, can significantly hinder accurate downstream characterization of many natural and designed protein systems. This result underscores the crucial role excipients played in eliminating DLST from our purification process while significantly improving the overall yield of the on-target assembled material. A comprehensive analysis of this on-target designed two-component nanoparticle will be detailed in an upcoming manuscript.

**Figure 3.**
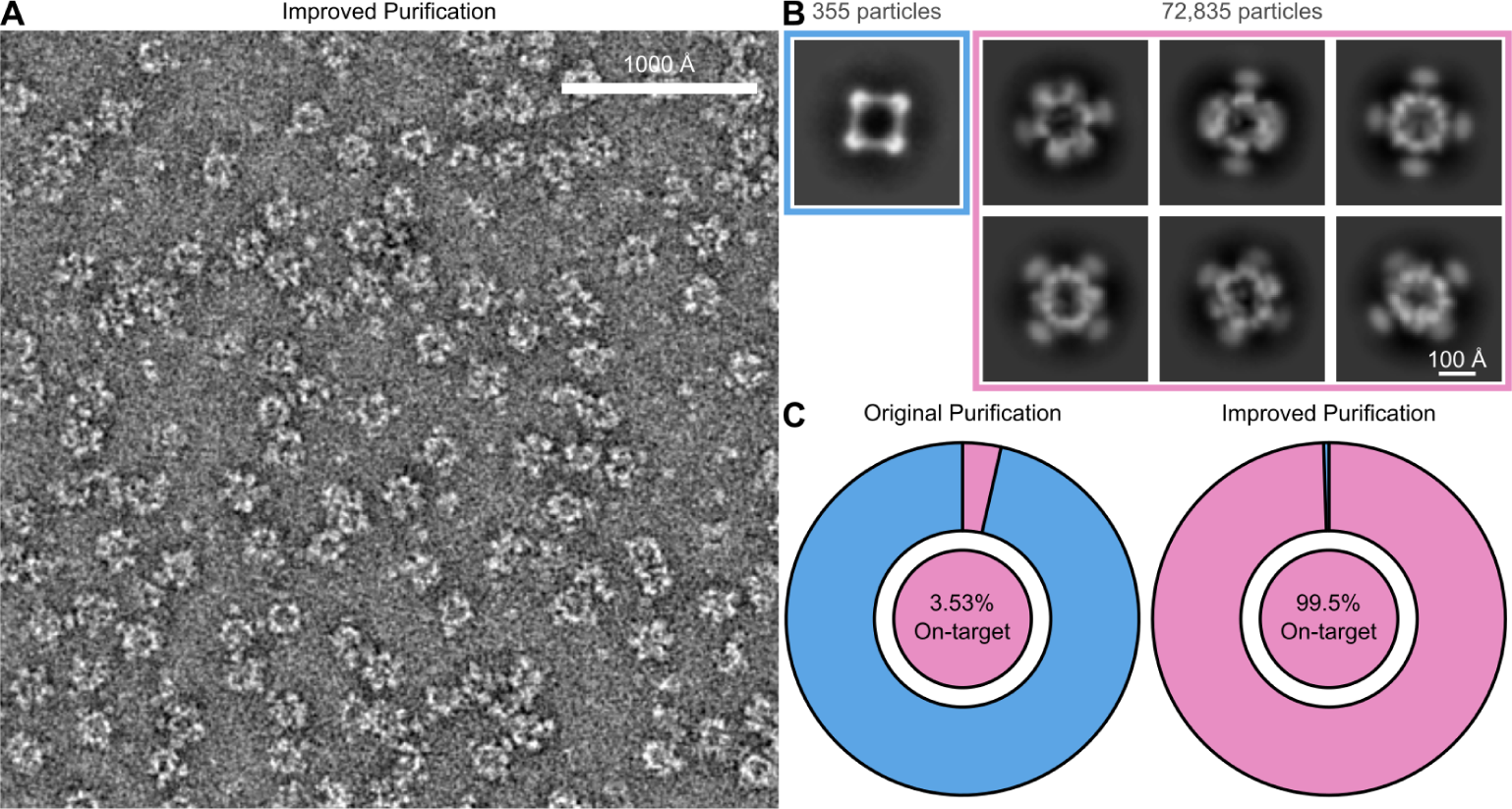
Modifications to the protein purification protocol result in an increased sample purity. (A) Representative ns-EM micrograph utilizing the optimized purification protocol. (B) Corresponding 2D ns-EM class averages of the on-target nanoparticle and DLST, with corresponding particle numbers listed for each. (C) Pie charts showing the relative abundance of the on-target nanoparticle and DLST as processed using the original purification protocol and the improved purification protocol. (Blue = off-target contaminant protein, DLST; Pink = on-target two component nanoparticle).

### Effect of Cryo-EM resolution on accurate DLST identification and potential broader applicability

The unambiguous identification of DLST as the protein contaminant using ModelAngelo and BLAST greatly facilitated our literature search and informed the rational refinement, iteration, and optimization of our protein purification protocol. Following this success, we aimed to explore the potential broader applicability of this ModelAngelo-to-BLAST identification approach, particularly under conditions where generating near-atomic resolution Cryo-EM data might not be feasible. Establishing the effectiveness of this workflow across a range of resolutions would increase its utility for various experimental setups, including those that often struggle to achieve high resolution. Specifically, we evaluated the ability of the pipeline to accurately identify DLST using Cryo-EM data spanning a broad resolution range. We first low-pass filtered the original 2.51 Å Cryo-EM map to resolutions of 3.0 Å, 4.0 Å, 4.5 Å, and 5.0 Å **(Figure 4A)**, which were then used as inputs for the ModelAngelo-to-BLAST pipeline. The average length of the sequence fragments generated by ModelAngelo was inversely proportional to the map resolution, with lower resolution input maps yielding shorter length chain fragments **(Figure 4B-C, Supplemental Table 4)**. Following the analysis of consensus sequences using Protein BLAST, the proportion of the top 100 results corresponding to DLST was plotted against their associated 3D map resolution values **(Figure 4D, Supplemental Sequences 1-5, Supplemental Tables 5-8)** [5]. This analysis revealed that at resolution thresholds of 2.51 Å, 3.0 Å, 4.0 Å, and 4.5 Å, BLAST accurately identified DLST as the most likely identity of the unknown protein in 98%, 99%, 99%, and 98% of cases, respectively **(Figure 4D)**. However, at 5.0 Å resolution, the consensus sequence did not yield any significant BLAST results, indicating a lack of detectable sequence similarity among known proteins for sequence fragments generated by ModelAngelo at this resolution **(Figure 4D)**. These results suggest that this pipeline could be applied to Cryo-EM data of other unknown contaminant proteins even when near-atomic resolution cannot be achieved. This flexibility across resolution ranges makes the method potentially more broadly applicable to different biological and experimental contexts, enabling researchers to confidently identify unknown proteins even in challenging or lower-resolution datasets.

**Figure 4.**
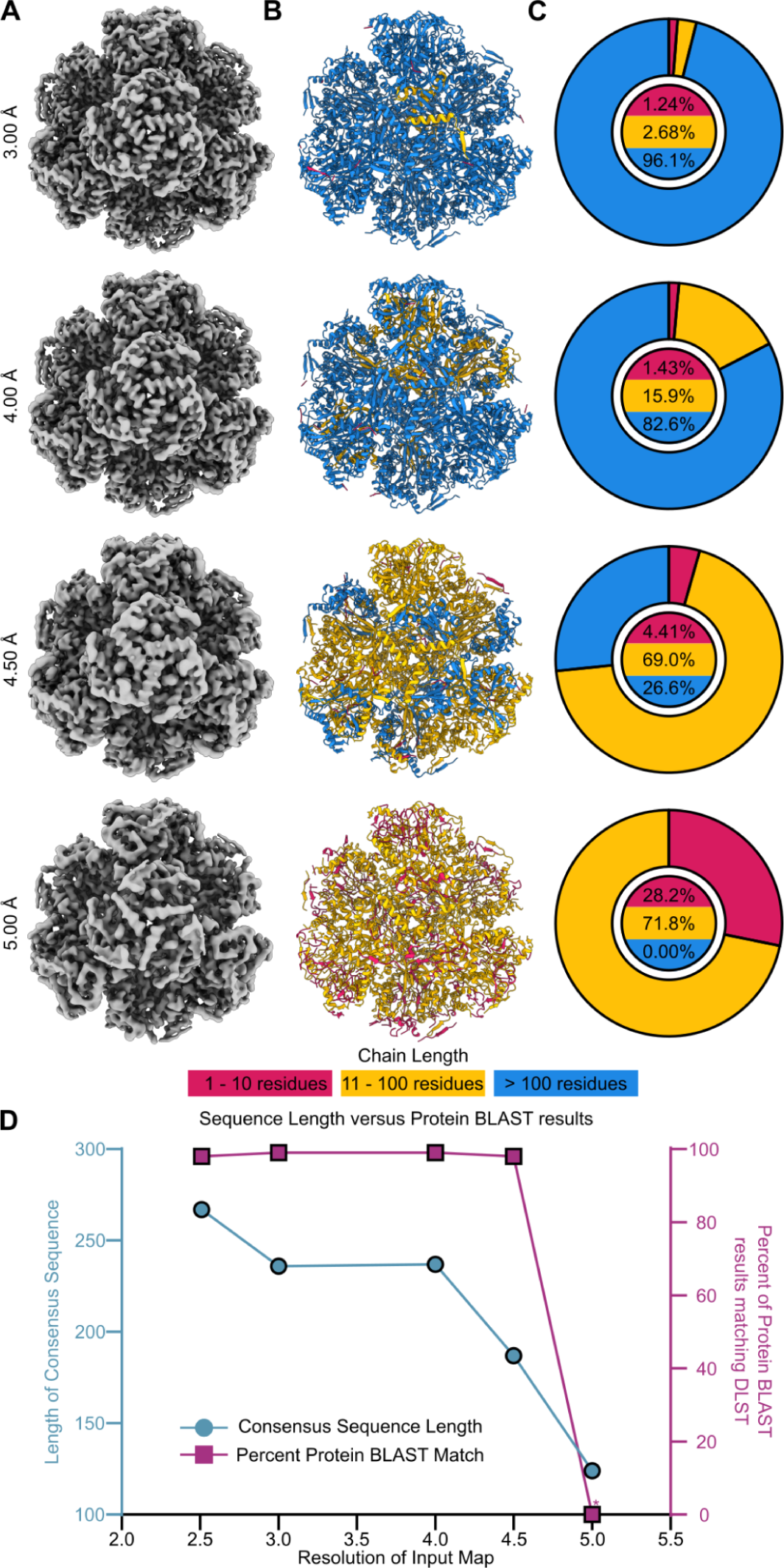
Impact of Cryo-EM resolution on accurate DLST identification. The Cryo-EM data was low pass filtered to 3.0 Å, 4.0 Å, 4.5 Å, and 5.0 Å and each map was used as the input for the computational steps of our workflow to evaluate the impact of resolution on the ability to identify DLST. (A) Low pass filtered Cryo-EM maps used for resolution-based analysis viewed along the 3-fold axis of symmetry. (B) sequence-agnostic ModelAngelo output with chains colored according to their lengths viewed along the 3 fold axis of symmetry. Chains with 1 - 10 residues are colored pink, chains with 11 - 100 residues are colored yellow, and chains longer than 100 are colored blue. (C) Pie charts of residues belonging to chains of a specified length as a percentage of all residues. Residues of a given chain length are colored according to the color of their chain as in B. (D) Three axis line plot showing the relationship between consensus sequence length and Protein BLAST results using input maps of different resolutions. At 5.00 Å the consensus sequence had a length of 124 residues and Protein BLAST was not able to generate results matched to its database.

## Discussion

Addressing the challenge of co-eluting protein contaminants during protein purification is essential for the production and accurate experimental characterization of recombinant proteins. In the case of the designed octahedral two-component nanoparticle system discussed here, the presence of a co-purifying protein artificially inflated the estimated concentration of the designed components required for nanoparticle assembly, leading to inconsistent assembly conditions and inaccurate estimations of the final on-target nanoparticle concentration **(Supplemental Table 1, Figure 1)**. Thus, it became necessary to identify and exclude this protein from our purifications so that we could more effectively produce and characterize our on-target design. By applying the ModelAngelo-to-BLAST pipeline and cross-referencing our cryoEM data with experimentally determined structures from the PDB, as well as AlphaFold 3 predicted structural information, we were able to unambiguously identify the co-purifying protein as DLST and optimized our purification protocols to remove it.

Before obtaining secondary structure information from Cryo-EM and implementing the ModelAngelo-to-BLAST pipeline, we hypothesized that the co-eluting protein might be the result of cross-sample contamination from another computationally designed protein purification. During our investigation, we explored several methods for potentially determining the identity of the unknown protein. One study utilizing Cryo-EM had successfully identified a number of unknown proteins found in native cell extracts utilizing the software package, Omokage [27]. However, this approach required existing structural information to confidently identify the unknown protein species [27,28]. During a retrospective analysis of our data, Omokage identified three computationally designed proteins (EMDB: 21163, 21164, 23266) among the top 10 potential sources of contamination, alongside DLST.

We also considered traditional experimental methods for unambiguous identification as well, such as liquid chromatography-tandem mass spectrometry (LC-MS/MS) and electrospray ionization mass spectrometry (ESI-MS). While these techniques are highly valuable, they posed significant anticipated challenges for our system, primarily due to our reliance on detergents to stabilize our designed two-component nanoparticle. Indeed, detergents are well-known to complicate mass spectrometry workflows and the interpretation of final data, adding significant complexity to an already complicated purification process [29]. Unlike the Omokage and mass spectrometry methods, the ModelAngelo-to-BLAST pipeline did not rely on prior PDB structural information, and enabled us to accurately identify the contaminant using only our experimental Cryo-EM data. Further validation using AlphaFold 3 models was as robust as PDB models during docking into the Cryo-EM density map, demonstrating that ModelAngelo-to-BLAST identifications can be verified even without the availability of pre-existing PDB information or manual building [19,28]. A key strength of this approach is its flexibility across different resolution ranges. While we initially used a near-atomic resolution cryo-EM map for the ModelAngelo-to-BLAST pipeline, we found that much lower resolutions were still sufficient to accurately identify DLST in our sample, aligning with previous benchmarking results for ModelAngelo [4]. This capability broadens the method’s applicability, making it practical for diagnosing issues in non-routine purifications as well as for exploratory studies where near-atomic resolution may be difficult to achieve.

Regarding the high-resolution cryo-EM structure of the DLST catalytic domain determined in this study, we note that minimal differences were observed between our structure and other previously reported crystallographic and cryo-EM structures of the DLST catalytic domain **(Supplemental Figure 5C-F)**. However, the resolution we achieved here did allow for the clear construction of a detailed intramolecular water network, which was particularly well-resolved at the catalytic pocket **(Supplemental Figure 5A-D)**. Comparison of this water network with the highest resolution published structure revealed key differences in the assignment of H_2_O atomic coordinates **(Supplemental Figure 5C-F)** [20]. These differences may reflect differences in the protein crystallization conditions versus the native-like environment of Cryo-EM sample preparation, potentially offering valuable insights for future studies aimed at understanding this enzyme’s mechanistic activity.

Ultimately, by leveraging advanced tools such as Cryo-EM, ModelAngelo, BLAST, the PDB, and AlphaFold 3, we transformed a challenging purification problem into an opportunity to refine our techniques and enhance the quality of our results. This integrated approach underscores the potential of combining cryo-EM with emerging AI and sequence-based tools as robust alternatives for tackling long-standing biochemical challenges—particularly in purifying proteins from complex biological systems, even when near-atomic resolution data cannot be achieved.

## Materials and Methods

### Protein purification of the two-component octahedral nanoparticle

The target two-component octahedral nanoparticle was designed *in silico* using a *de novo* designed tetramer and a soluble trimer based on the aldolase 2-keto-3-deoxy-6-phosphogluconate (KDPG). Specifics on the components for this designed system will be discussed further in a separate manuscript detailing this design. Thousands of octahedral nanoparticle designs were screened *in silico* through careful evaluation of the interface residues. Filters were applied to all designs during *in silico* screening until the top 10 nanoparticles were chosen to be evaluated *in vitro*. For the top designs, C-terminal histidine-tagged plasmids were ordered from GenScript and transformed into *E. coli* BL21 DE3 competent cells.

For the designed KDPG trimer, *E. coli* were grown in Terrific Broth media at 37.0°C until an OD600 of 0.60 - 0.80 before induction with Isopropyl β- d-1-thiogalactopyranoside (IPTG) and grown for an additional 3 hours at 37.0°C. Cells were harvested at 5000 rpm for 15 minutes and resuspended with the lysis buffer (25.0 mM Tris-HCl pH 8.00, 0.15 M NaCl, 100 mM arginine, 100 mM glycine, 1.00 mg/mL lysozyme and 1.00 mg/mL DNAse) then lysed using sonication. Lysed cell resuspensions were sonicated for a total of 5 minutes using 10.0s sonication pulses and 10.0s breaks. After cell lysis, the solution was pelleted at 10,000*g* for 30 minutes. The supernatant was collected and ran through a nickel-nitrilotriacetic acid (Ni-NTA) IMAC resin with a bed volume of 5.00 mL. The IMAC resin was equilibrated with 5.00 column volumes of loading buffer (25.0 mM Tris-HCl pH 8.00, 0.15 M NaCl, 100 mM arginine and 100 mM glycine) before the addition of cell lysate, then washed with 3 column volumes of the loading buffer. The bound protein was eluted with 1.50 column volumes of elution buffer (25.0 mM Tris-HCl pH 8.00, 0.15 mM KCl, 0.50 M imidazole, 0.15% n-Dodecyl-B-D-maltoside (DDM), 100 mM arginine and 100 mM glycine).

For the tetrameric protein component, plasmids were transformed into BL21 Star (DE3)pLysS One Shot competent cells. Cells were grown for 3 hours at 37.0°C until an OD600 of 0.80 - 1.00 before induction with Isopropyl β- d-1-thiogalactopyranoside (IPTG) and grown for an additional 18 hours at 18.0°C. The cells were then lysed in the same fashion as described above. Following lysis by sonication, cells expressing the tetramer were spun at 10,000*g* for 15 minutes and the supernatant was collected for ultracentrifugation at 130,000*g* for 1.00 hour at 4.0°C. A resuspension buffer (25.0 mM Tris-HCl pH 8.00, 0.15 M KCl, 100 mM arginine, 100 mM glycine and 2.00% DDM) was used to resuspend the pellet after ultracentrifugation. This resuspension was placed on a rocker at 4.0°C overnight. The following day, the resuspended solution was spun for an additional 30 minutes at 14,000*g*. The supernatant was collected and ran through an Ni-NTA IMAC resin with a bed volume of 5.00 mL. The IMAC resin was equilibrated with 5.00 column volumes of resuspension buffer before the addition of cell lysate, then washed with 3 column volumes of the resuspension buffer and eluted with 1.50 column volumes of elution buffer (25.0 mM Tris-HCl pH 8.00, 0.15 mM KCl, 0.50 M imidazole, 0.15% n-Dodecyl-B-D-maltoside (DDM), 100 mM arginine and 100 mM glycine).

Purified proteins were mixed together at a 1:1 (c/c) by molar ratio for 1 hour at 4.0°C to facilitate nanoparticle assembly. Further purification of the nanoparticles was achieved using size exclusion chromatography (SEC) in an SEC buffer (25.0 mM Tris-HCl pH 8.00, 0.15 mM NaCl, 0.10% DDM, 100 mM arginine and 100 mM glycine) on a Cytiva Superose® 6 Increase 10/300 GL column. The SEC column was calibrated using the Cytiva Gel Filtration HMW Calibration Kit (CAT 28403842) with molecular masses ranging from 43,000 to 669,000 Daltons. The nanoparticle peak purified on the SEC was isolated and characterized by DLS, ns-EM and Cryo-EM.

All lysis, resuspension and purification buffers resulting in the co-purification of DLST with the on-target nanoparticle did not contain 100 mM glycine and 100 mM arginine. Otherwise, the purification protocols described here are the same for samples where co-purification of DLST with our two-component nanoparticle were observed.

### ns-EM sample preparation

After SEC, the purified peak was diluted from 0.12 mg/mL to 0.10 mg/mL using the SEC buffer. 3 µL of the diluted sample was applied to a freshly glow discharged 400 carbon square mesh grid (Electron Microscopy Sciences, EMS) and allowed to adsorb onto the grid for 30 seconds. Blotting paper (Wattman) was used to remove excess solution from the grid before the application of 3 µL 2% uranyl formate (2UF) to stain the grid for ns-EM. Excess stain solution was blotted immediately followed by another application of 3 µL 2UF. This was repeated once more, resulting in a total of 3 applications of 2% UF. Following the final application of 2% UF, all remaining solution was blotted and the sample was allowed to dry completely. There was no waiting period in between each application of 2UF, and after each blot new 2UF was applied to the grid immediately.

### ns-EM data acquisition and processing

Negative stain electron microscopy (ns-EM) micrographs were collected on a Thermo Fisher Scientific Talos L120C 120 kV transmission electron microscope with a LaB_6_ filament and CETA camera with a pixel size of 2.49 Å at 57,000x magnification. 164 micrographs were recorded at 57,000x magnification at a total dose rate 40.8 e^-^/Å^2^ for the octahedral nanoparticle. Micrographs were imported into CryoSPARC v4.4 [9] and Patch CTF Estimation was performed prior to blob picking, inspection, and extraction of 22,158 picks with a box size of 180 pixels (448.2 Å). With CTF correction off, picks were classified into 50 2D classes, four of which were selected to serve as templates for an additional round of particle picking. In this second round of particle picking, picks were inspected and extracted to 180 pixels (448.2 Å) before classification of 55,802 extracted particles into 150 2D classes with CTF correction off. 5 out of 150 of these 2D classes were selected as a second round of templates for another round of particle picking and extraction. In this last round, 49,262 particles were extracted to 180 pixels (448.2 Å) using the templates from the second round of extraction and classified into 50 2D classes with a batch size of 400 particles and CTF correction off. 30,927 particles were used to homogeneously refine a map into Octahedral symmetry with a maximum resolution threshold of 20 Å. Particles were aligned into three 3D octahedral ab-initio reconstruction classes. For subsequent ns-EM data processing for DLST, we followed a similar pipeline outlined here.

### Cryo-EM sample preparation

Prior to freezing, 3 µL of protein sample, at an estimated final concentration of 0.3 mg/mL, was applied to glow-discharged Quantifoil 2/2 holey carbon grids overlaid with a thin layer of additional carbon. Vitrification was performed using a Vitrobot MkIV at 22°C and 100% humidity, with a wait time of 7.5 seconds, a blot time of 0.5 seconds, and a blot force of 0, before being immediately plunge-frozen into liquid ethane. The sample grids were clipped following standard protocols before being loaded into a 300 kV Titan Krios for imaging.

### Cryo-EM data acquisition and processing

A total of 6,211 movies were collected using SerialEM [30,31] on a 300 kV FEI Titan Krios equipped with a Gatan K3 direct electron detector and an energy filter. The movies were recorded at 105,000x magnification, with a pixel size of 0.843 Å/pixel. Each movie consisted of 99 frames recorded at a frame rate of 19.8 frames per second, with a dose rate of 9.4 e-/Å²/s and an exposure time of 5 seconds, resulting in a total exposure dose of 47.0 e-/Å². All movies were imported into CryoSPARC v4.4 for data processing [9]. Initially, the movies were corrected for Patch Motion Correction. Defocus and astigmatism values were estimated using the Patch CTF with the default parameters. After two rounds of exposure curation, 4,264 exposures were selected for particle picking. Prospective particles were identified using blob picking with a diameter range of 200 to 400 Å, corresponding to the expected sizes of both our intended and co-purified proteins. This process identified 808,774 prospective particles, which were then inspected and curated based on power histogram values and normalized cross-correlation scores. Following this curation, 456,319 particles were extracted at a box size of 600 pixels. These particles were classified into 150 classes with a batch size of 400 per class. 4 classes were selected to serve as templates for another round of particle picking with particle diameter set to 300 Å. This round yielded 1,468,115 prospective particles, which were curated down to 765,699 particles. These were extracted at 600 pixels and classified into 150 classes again. All high-quality particle classes (47,980 particles) were re-extracted with recentering enabled, resulting in 47,480 particles, which were then split into 150 classes, yielding 39,222 high-quality particles. Of these, 19,033 of the unknown protein contaminant particles were reconstructed into 3 ab-initio 3D C1 reconstruction classes. All particles were homogeneously refined into the best Ab-initio volume with octahedral symmetry enabled, achieving a resolution of 2.62 Å. Local CTF refinement was performed, followed by another round of homogeneous refinement, improving the resolution to 2.59 Å. Reference Based Motion Correction was then performed which reduced the number of particles to 18,809. Subsequently, Global CTF refinement was run for two iterations with Fit Anisotropic Mag set to true. One final homogenous refinement was run in Octahedral symmetry with Minimize over per-particle scale and Optimize per-group CTF parameters set to true, yielding a map with a final estimated resolution of 2.51 Å.

### Identification of co-purified protein contaminant from octahedral nanoparticle samples

ModelAngelo was run on the final Cryo-EM map, sharpened using deepEMhancer, in a sequence-agnostic manner to generate sequence segments fit to the density as a cif file [34]. PyMol [32] was used to generate a FASTA sequence [11] from the cif file which was aligned by the multiple sequence alignment program Clustal Omega [10]. The aligned sequences were then exported and used by sequenced logo program WebLogo [33] to generate a frequency plot of the aligned sequences. The most frequent amino acid present at each residue was used to create a consensus sequence. This sequence was input into the National Institutes of Health (NIH)’s Protein BLAST [5] and the result strongly indicated that the most likely candidate protein was dihydrolipoamide succinyltransferase **(Table 3)**. PDB assembly 1SCZ [20] of the catalytic domain of dihydrolipoamide succinyltransferase was aligned to our Cryo-EM volume and used to visually confirm the identity of the contaminant.

### Cryo-EM Structural Refinement

Following identification of the contaminant protein as DLST, the *dlst* gene sequence from BL21 DE3 *E. coli* was pulled from UniProt (entry A0A140NDX4) [16,17] and used as the input sequence for automated model building using ModelAngelo along with the 2.51 Å deepEMhancer [34] sharpened volume map. The resulting model was imported into Pymol [32] to extract its FASTA [11] sequence, the chains of which were then aligned with the UniProt gene sequence once more to verify that deviations to the sequence were not introduced. Then the model was relaxed using ISOLDE [35] in ChimeraX [36–38] to ensure the model fit moderately well to the density. Subsequently, one chain of the model was isolated and retained while all other copies were deleted. Symmetry operators were applied to the remaining chain in ChimeraX [36–38] to repopulate the density with the 23 additional subunits. The model was then iteratively processed using Coot [39], Phenix [40,41], and ISOLDE [35] to accurately orient the backbone and sidechains. After each iteration of processing, the A chain was preserved and other chains were first deleted, and then repopulated using ChimeraX [36–38] symmetry operators. Periodically, between iterations, the model was submitted to the wwPDB validation service [42] to provide an additional metric of model quality. Once the iteratively produced model was deemed by the authors to be of sufficient quality, a mask was generated in ChimeraX [36–38] of the 2.51 Å map sharpened (without using deepEMhancer [34]) preserving only regions within 6 Å of chain A. A network of waters with density present in this mask was built in Coot [39] and then symmetrized over the whole model in ChimeraX [36–38]. The symmetrized model was examined in ChimeraX and Coot to identify regions where waters were duplicated (as waters were built along all interface betweens chain A and the adjacent chains), all but one copy of each of these waters were deleted such that following an additional round of symmetrization the final model possessed no duplicates. The full model was refined one last time in Phenix [40] using the 2.51 Å map sharpened (without using deepEMhancer) map. The distance between each water and nearby non-Hydrogen atoms was measured to verify that the waters were not placed too near or far from the model. The model was manually inspected in Chimera [43] as a final quality check and then the final model was once again submitted to the wwPDB validation service [42] to provide a quantitative assessment of model quality prior to deposition.

## Acknowledgements

We would first like to thank the developers of ModelAngelo, without whom we would probably still be banging our heads against the bench, desperately cleaning SEC columns in an attempt to remove the contaminant from our *de novo* designed protein samples. We would also like to thank Rebecca Cole and Rashmi Ravichandran for their efforts in protein expression, purification, and evaluation, and Joel Quispe and Sasha Dickinson for management of the University of Washington Cryo-EM facilities. Additionally, we thank Kandise VanWormer, Hernan Nunez-Ortega, Rafael Ticzon, Andre Dubief, and Garrett Ruth for providing laboratory support as well as Luki Goldschmidt, Patrick Vecchiato, and Bulat Faezov for technical support. Molecular graphics and analyses performed with UCSF ChimeraX, developed by the Resource for Biocomputing, Visualization, and Informatics at the University of California, San Francisco, with support from National Institutes of Health R01-GM129325 and the Office of Cyber Infrastructure and Computational Biology, National Institute of Allergy and Infectious Diseases. Pie Charts used in figures were prepared using GraphPad Prism version 10.1.0 for Windows, GraphPad Software, Boston, Massachusetts USA, www.graphpad.com. The graphical abstract was created with Biorender.com.

## Author Contributions

K.D.C., D.Z., and A.J.B. conceptualized the research. D.Z. designed the *de novo* two-component nanoparticle primarily discussed in this manuscript and expressed and purified the original two-component nanoparticle samples containing DLST. K.D.C. and D.Z. performed ns-EM. A.G., H.E.E., H.P., and A.C. contributed additional ns-EM micrographs. A.J.B. prepared samples for Cryo-EM. C.W. and A.J.B conducted Cryo-EM sample screening and data collection. K.D.C. processed the Cryo-EM data. K.D.C. outlined, tested, and ran the ModelAngelo-to-BLAST pipeline for the accurate identification of DLST. K.D.C. and C.W. built the DLST Cryo-EM structure. D.Z. optimized the purification protocol to exclude DLST. K.D.C. performed resolution-based analyses to benchmark the ModelAngelo-to-BLAST protein identification pipeline against lower resolution Cryo-EM data. N.P.K. and A.J.B. provided supervision. K.D.C., D.Z., C.W., and A.J.B. wrote the manuscript with input from all authors.

## Supplemental Figures

**Supplemental Figure 1.**
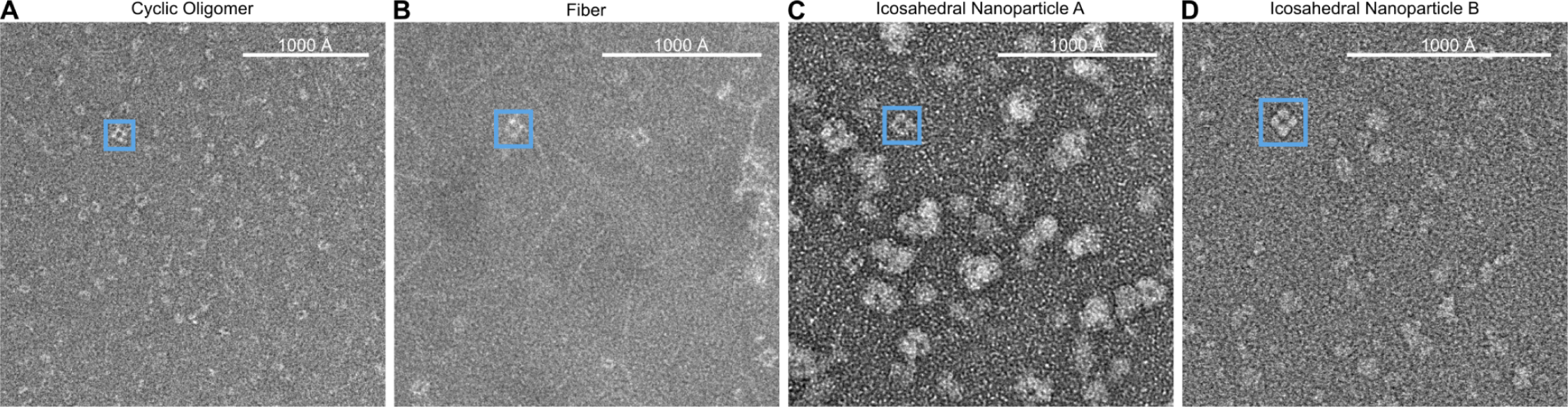
Octahedral contaminant protein observed in additional de novo protein samples by negative stain electron microscopy. Sections of representative ns-EM micrographs demonstrating observation of the octahedral contaminant protein in multiple samples of *de novo* proteins imaged at various magnifications, with 1000 Å scale bars sized proportionally. (A) A sample of a *de novo* cyclic-oligomeric protein imaged at 45,000x magnification. (B) A sample of a *de novo* protein fiber imaged at 57,000x magnification. (C) A sample of an icosahedral nanoparticle imaged at 57,000x magnification. (D) A sample of a different *de novo* icosahedral nanoparticle imaged at 73,000x magnification.

**Supplemental Figure 2.**
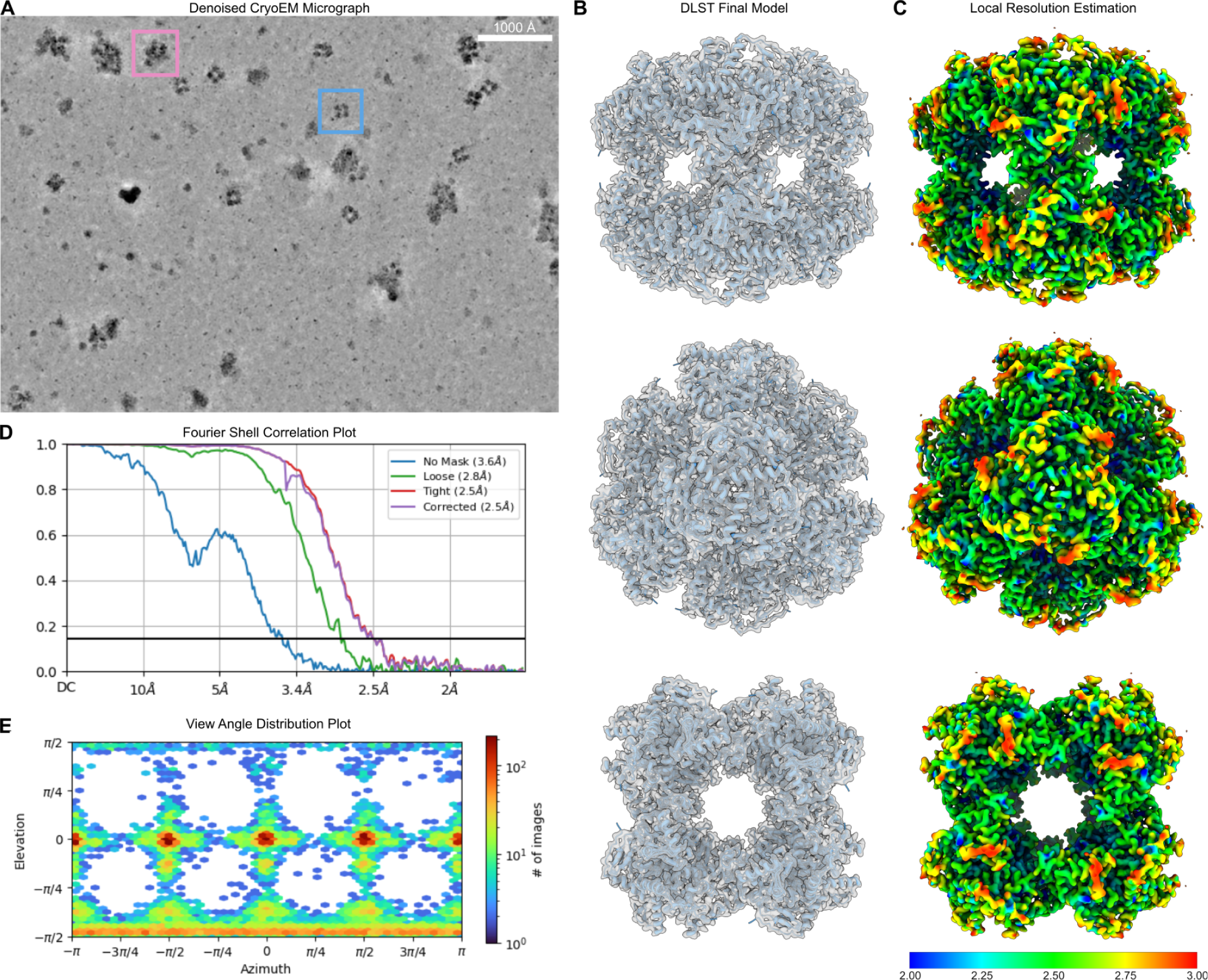
Cryo-EM Data processing metrics. (A) Denoised representative Cryo-EM micrograph at 105,000x magnification showing particles of both the designed two-component nanoparticle and DLST. (Blue = off-target contaminant protein; Pink = on-target two component nanoparticle). (B) The final built model of the DLST catalytic domain docked into the 2.51 Å density map along the 2, 3, and 4 fold axes of symmetry. (C) Local resolution estimate of the Cryo-EM map with a 0.143 FSC cutoff. Colors range from blue at 2.00 Å to red at 3.00 Å. (D) Fourier shell correlation plot of the Cryo-EM map. (E) View angle distribution plot of the Cryo-EM map.

**Supplemental Figure 3.**
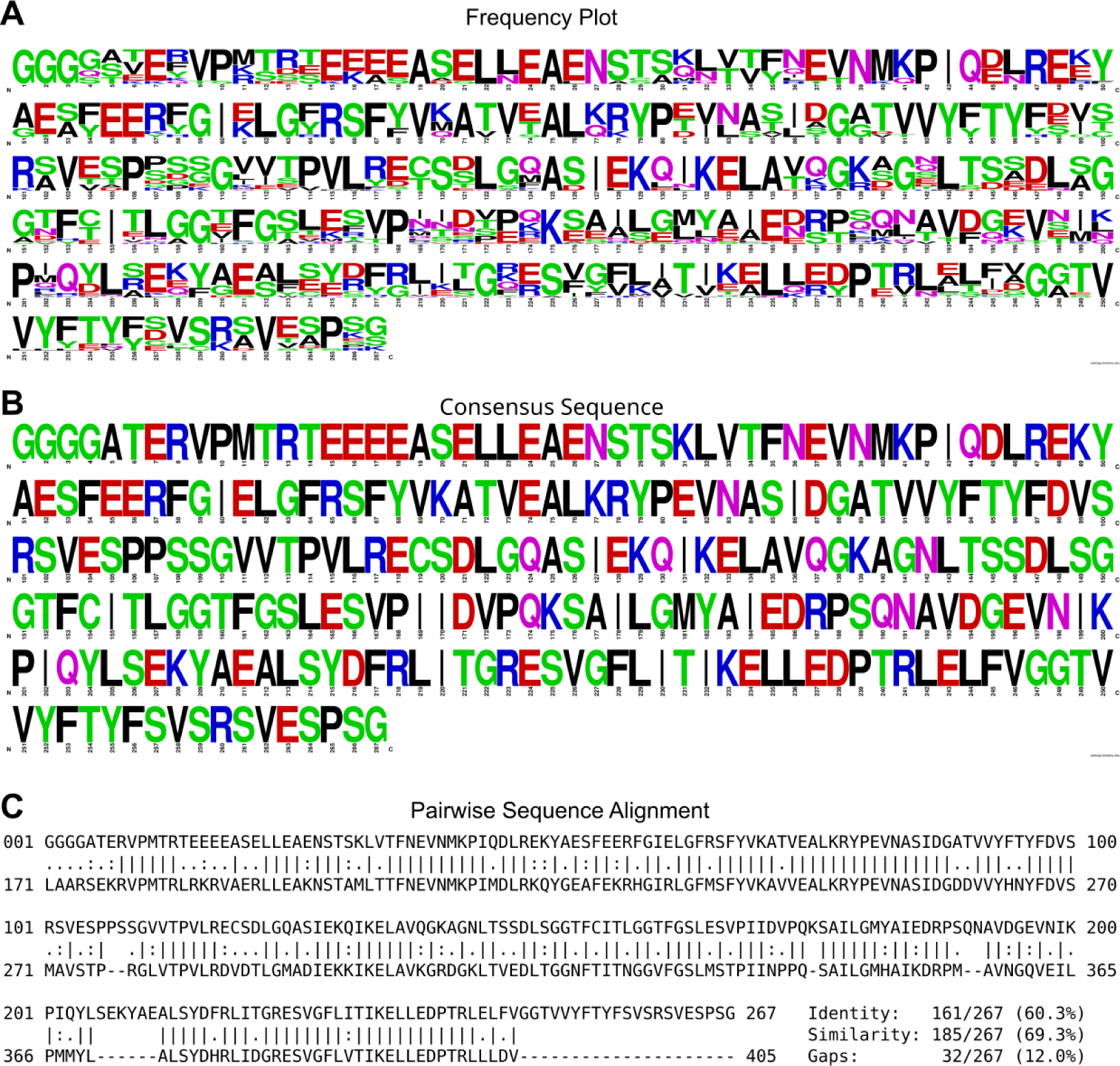
Sequences derived from ModelAngelo’s sequence-agnostic method used to identify and model DLST from the Cryo-EM volume map. (A) ModelAngelo sequence fragments were aligned in an MSA to generate a frequency plot displaying relative abundance of amino acids at each position. (B) A consensus sequence for one chain was generated using amino acid frequency data from (a) at each position. This consensus sequence was the input for protein BLAST [5]. (C) After protein BLAST identified DLST, UniProt was searched for the corresponding DLST sequence for the BL21 (DE3) expression vector. The UniProt sequence A0A140NDX4 (bottom) [17] was aligned with the consensus sequence (top). The first alignment revealed a gap spanning the first 170 residues of the UniProt sequence so an additional alignment was performed using only the UniProt residues at position 170 or later.

**Supplemental Figure 4.**
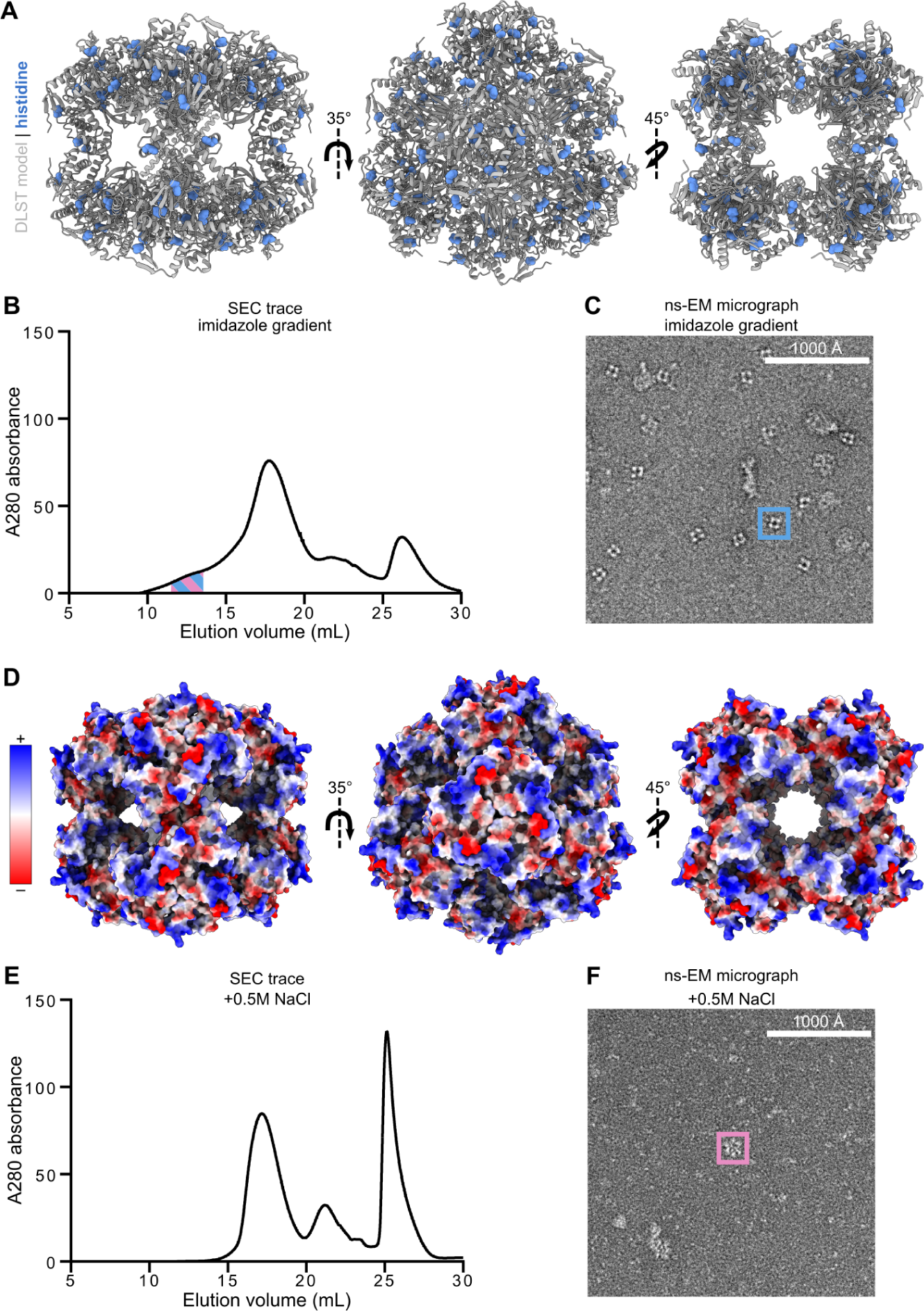
Improving purification of DLST through modification of purification protocols. Various purification attempts of the on-target nanoparticle were conducted, evaluating buffer conditions to eliminate the co-purification of DLST. (A) Location of histidines (blue) in DLST along the 2, 3, and 4 fold axis of symmetry. (B) SEC trace of the octahedral nanoparticle after protein components were purified with an elution gradient from 0.00 M - 0.50 M imidazole. (C) One fraction from SEC purification was taken for ns-EM. This is a cropped view of the ns-EM micrograph taken at 57,000x magnification. The majority of observed particles were of DLST. (D) Electrostatic potential of the catalytic domain of DLST with positively charged residues shown in blue and negatively charged residues in red along the 2, 3, and 4 fold axes of symmetry. (E) SEC trace of the octahedral nanoparticle after cells were lysed with 0.5 M NaCl. IMAC buffers to purify components also contained 0.5 M NaCl. (F) One fraction from SEC where the expected elution time of the nanoparticle was taken for ns-EM. This is a cropped view of the ns-EM micrograph taken at 57,000x magnification. Particle total yield and total density of assembled material was significantly diminished. We did not observe the presence of DLST in these micrographs. (Blue = off-target contaminant protein, DLST; Pink = on-target two component nanoparticle).

**Supplemental Figure 5.**
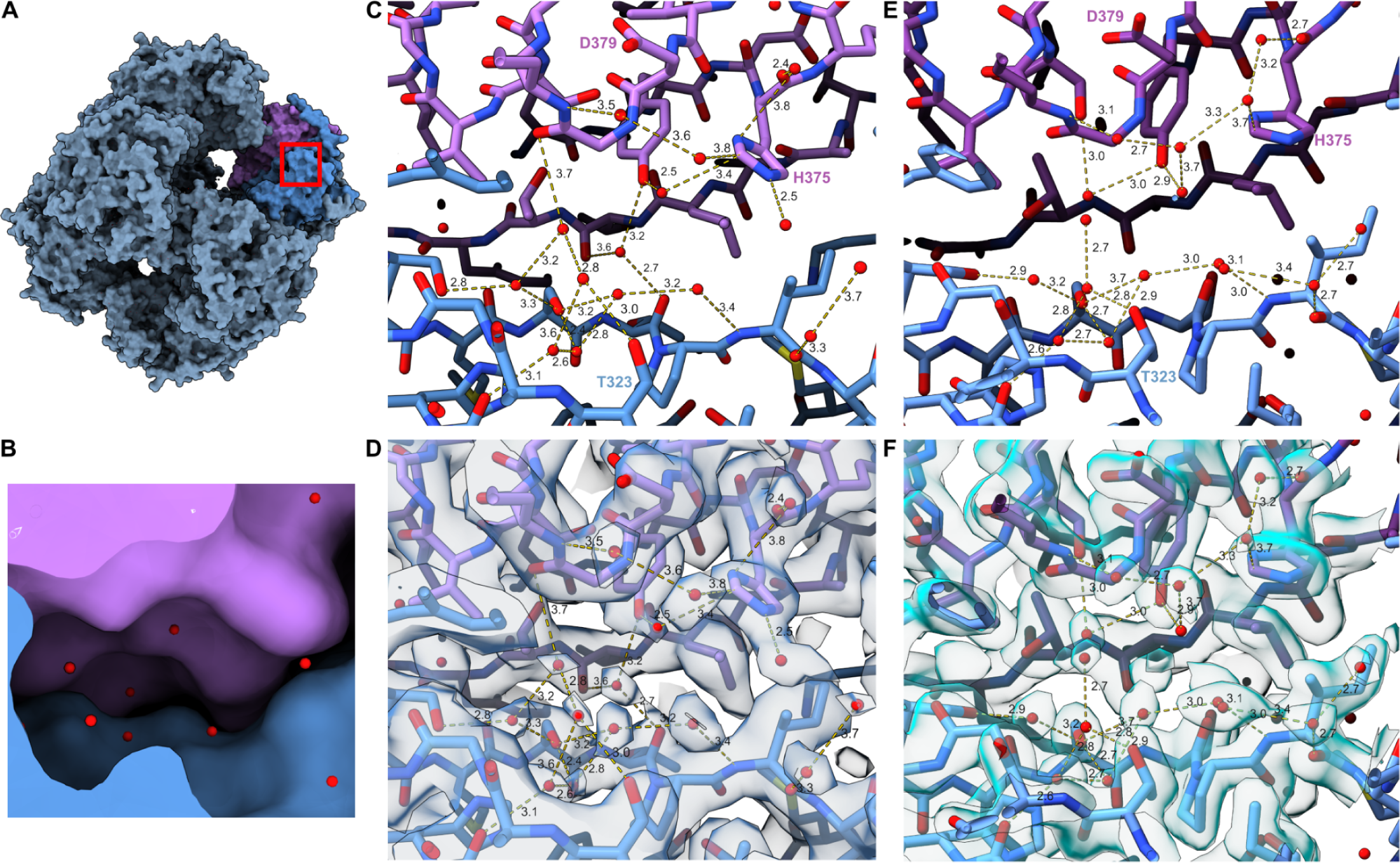
A comparison of the active site water network coordinates for DLST. Water network of the DLST active site comparing the 2.51 Å Cryo-EM structure discussed here with the 2.20 Å crystal structure (PDB: 1SCZ) [20]. The catalytic residues HIS375, ASP379, THR323 are labeled. (A) Our Cryo-EM structure surface view, with a red box showing the location of the active site. (B) A zoomed in view of the active site highlighted in (a) using a cutaway surface view. (C) The Cryo-EM structure of DLST built from the 2.51 Å Cryo-EM map. (D) The structure in (c) fit within the Cryo-EM map. (E) 1SCZ model taken from the deposited structure [20]. (F) 1SCZ fit within the 2Fo-Fc map [20].

**Supplemental Table 1.**
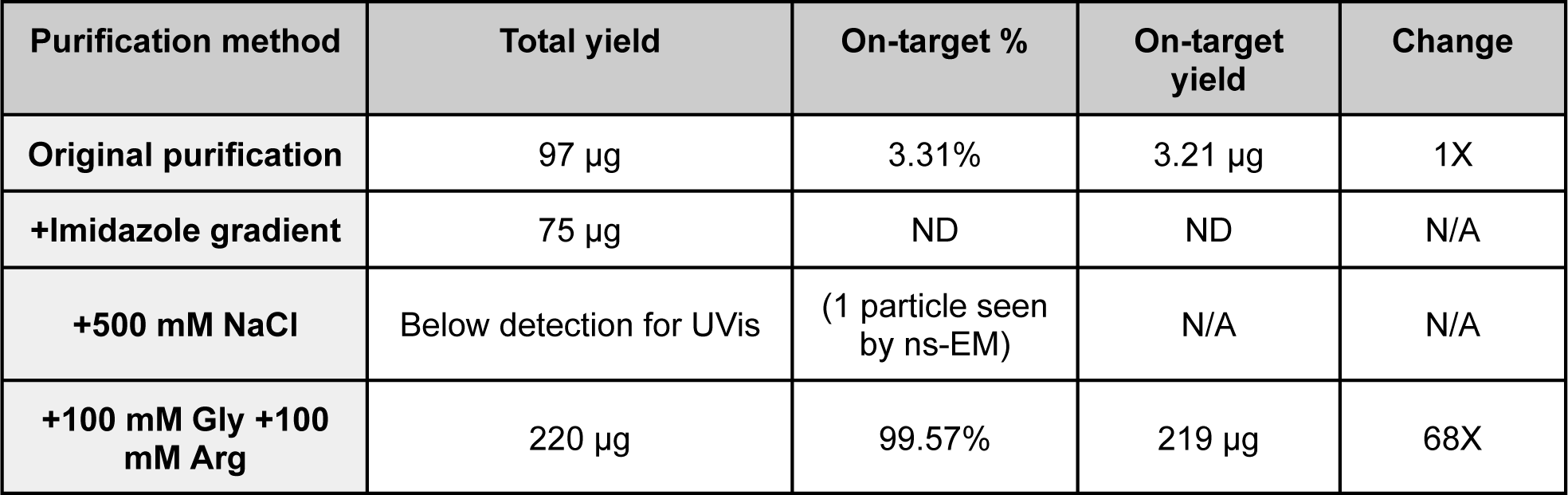
Protein yields from purification methods.

**Supplemental Table 2.**
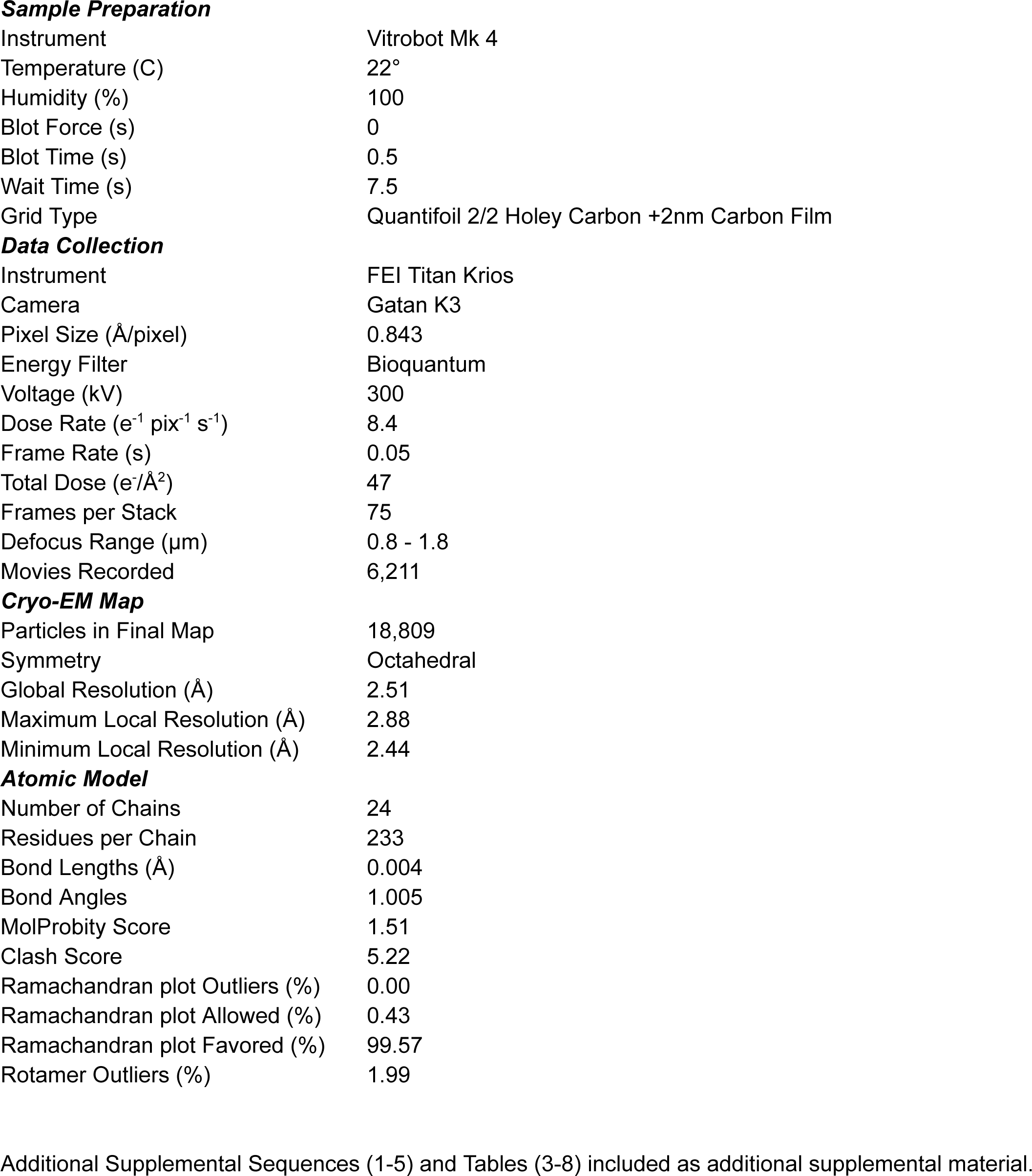
Cryo-EM Data Collection, Map, and Model Statistics.

**Sample Preparation**
|  |  |
| --- | --- |
| Instrument | Vitrobot Mk 4 |
| Temperature (C) | 22° |
| Humidity (%) | 100 |
| Blot Force (s) | 0 |
| Blot Time (s) | 0.5 |
| Wait Time (s) | 7.5 |
| Grid Type | Quantifoil 2/2 Holey Carbon +2nm Carbon Film |
| **Data Collection** |  |
| Instrument | FEI Titan Krios |
| Camera | Gatan K3 |
| Pixel Size (Å/pixel) | 0.843 |
| Energy Filter | Bioquantum |
| Voltage (kV) | 300 |
| Dose Rate (e<sup>-1</sup> pix<sup>-1</sup> s<sup>-1</sup>) | 8.4 |
| Frame Rate (s) | 0.05 |
| Total Dose (e<sup>-</sup>/Å<sup>2</sup>) | 47 |
| Frames per Stack | 75 |
| Defocus Range (µm) | 0.8 - 1.8 |
| Movies Recorded | 6,211 |
| **Cryo-EM Map** |  |
| Particles in Final Map | 18,809 |
| Symmetry | Octahedral |
| Global Resolution (Å) | 2.51 |
| Maximum Local Resolution (Å) | 2.88 |
| Minimum Local Resolution (Å) | 2.44 |
| **Atomic Model** |  |
| Number of Chains | 24 |
| Residues per Chain | 233 |
| Bond Lengths (Å) | 0.004 |
| Bond Angles | 1.005 |
| MolProbity Score | 1.51 |
| Clash Score | 5.22 |
| Ramachandran plot Outliers (%) | 0.00 |
| Ramachandran plot Allowed (%) | 0.43 |
| Ramachandran plot Favored (%) | 99.57 |
| Rotamer Outliers (%) | 1.99 |

Additional Supplemental Sequences (1-5) and Tables (3-8) included as additional supplemental material.

