## Additional Supplemental Data for "Protein identification using cryo-EM and artificial intelligence guides improved sample purification"

#### Sequence 1 | 2.51 Å sharpened volume map consensus sequence

GGGGATERVPMTRTEEEEASELLEAENSTSKLVTFNEVNMKPIQDLREKY
AESFEERFGIELGFRSFYVKATVEALKRYPEVNASIDGATVVYFTYFDVS
RSVESPPSSGVVTPVLRECSDLGQASIEKQIKELAVQGKAGNLTSSDLSG
GTFCITLGGTFGSLESVPIIDVPQKSAILGMYAIEDRPSQNAVDGEVNIK
PIQYLSEKYAEALSYDFRLITGRESVGFLITIKELLEDPTRLELFVGGTV
VYFTYFSVSRSVESPSG

#### Sequence 2 | 3.00 Å sharpened volume map consensus sequence

SGSEKRVPRSKERKKEADRLNEAKNSTAMLTTFQEVNMKPIQDLRKKYGE
AFEKRYGIRLGFRSFYVKAVVEALKRYPEVLASIEGDNVVYYNYFEVSRA
VETPSKGGLVTPVLREVSTLGMADIEKKIKELATKARDGKLTTNELEGGN
FTITNGGTFGSLMSVPIINPDPRAAILGLYAIKDRPQAVNGQVEILPMQY
LSLSYDFRLIDGRESVGFLVTIKELLEDPTRLLLFV

#### Sequence 3 | 4.00 Å sharpened volume map consensus sequence

GRSEERVPRRSKERKKVAERLNEALNSTVMIVVFNSVNMKPIQDLRKKYA
EAFEKRYGVRLGFRSYYVKAVVEALKRYPDVNASLEGDTVVYYNYFEVTK
AVETRKDKGLVTPCLREVSTLGMADIEKKIKELAVRARSGKLTEEELSGG
NFTITNSGEFGGALMSVPIINPPQAAILGMHSIKDRPQAVNGKVEILPKM
YLSLSYDYRIISAREATGFLTTIKELLEDPTRLLLFL

#### Sequence 4 | 4.50 Å sharpened volume map consensus sequence

SGSTERVPLRSKERKKLAERLNEALNSTSMTTVFVEVNMKPIQELRKKYA
EAFEKRRYGVRIGHRAFHIQAVVEALKRHPEIMASLEGDTVVYYTEFGVL
MSKPVDNPNCAGLVTLCLRSIKDLPMAINGQVELLPQMYLALSYDYRIIT
AREAVGFLLKIKKLAEKAREGDITREILSGAGDTPKK

#### Sequence 5 | 5.00 Å sharpened volume map consensus sequence

IDIRAKGTESVPIQEKRKKEAEAFEKAYGIKILGLRAFRLSLPVPGPVNR
YPELLGAYSPKTRPKANNGKVEVLPCSWLELEYSRRVVEARKLAEGFLEE
VKRELAEVARSGNITEELLLGGGA

Supplemental Table 3 | Designed v. Contaminant particles in samples using the original and modified purification protocols

| Purification Method | Proteins matching the designed nanoparticle | Contaminant proteins | Total proteins | Percent proteins matching the designed nanoparticle | Percent contaminant proteins |
|---|---|---|---|---|---|
| **Original Purification Protocol** | 1131 | 30927 | 32058 | 3.53% | 96.5% |
| **Modified Purification Protocol** | 72835 | 355 | 73190 | 99.5% | 0.49% |

Supplemental Table 4 | Length of chain fragments output by Model Angelo using various resolution input maps [Results limited to *E. coli* (taxid: 562)]

| Resolution of Map and version of ModelAngelo | Length of Chain | Quantity of Chains | Percent of Chains |
|---|---|---|---|
| **2.51 Å v1.0.4** | 001 - 010 | 50 | 54.35 |
| | 011 - 050 | 0 | 0.00 |
| | 051 - 100 | 1 | 1.09 |
| | 101 - 150 | 35 | 38.04 |
| | 151 - 200 | 0 | 0.00 |
| | 201 - 250 | 6 | 6.52 |
| **3.00 Å v1.0.4** | 001 - 010 | 51 | 64.56 |
| | 011 - 050 | 0 | 0.00 |
| | 051 - 100 | 2 | 2.53 |
| | 101 - 150 | 5 | 6.33 |
| | 151 - 200 | 1 | 1.27 |
| | 201 - 250 | 20 | 25.32 |
| **4.00 Å v1.0.4** | 001 - 010 | 52 | 57.14 |
| | 011 - 050 | 1 | 1.10 |
| | 051 - 100 | 13 | 14.29 |
| | 101 - 150 | 5 | 5.49 |
| | 151 - 200 | 9 | 9.89 |
| | 201 - 250 | 11 | 12.09 |
| **4.50 Å v1.0.4** | 001 - 010 | 131 | 57.46 |
| | 011 - 050 | 45 | 19.74 |
| | 051 - 100 | 40 | 17.54 |
| | 101 - 150 | 12 | 5.26 |
| | 151 - 200 | 0 | 0.00 |
| | 201 - 250 | 0 | 0.00 |
| **5.00 Å v1.0.4** | 001 - 010 | 471 | 74.41 |
| | 011 - 050 | 149 | 23.54 |
| | 051 - 100 | 13 | 2.05 |
| | 101 - 150 | 0 | 0.00 |
| | 151 - 200 | 0 | 0.00 |
| | 201 - 250 | 0 | 0.00 |
| **2.51 Å v1.0.12** | 001 - 010 | 49 | 55.06 |
| | 011 - 050 | 3 | 3.37 |
| | 051 - 100 | 3 | 3.37 |

| | | | |
|---|---|---|---|
| | 101 - 150 | 24 | 26.97 |
| | 151 - 200 | 0 | 0.00 |
| | 201 - 250 | 10 | 11.24 |

Supplemental Table 5 | Protein BLAST results based on consensus sequence of automated model building on a 2.51 Å sharpened volume map [Results limited to *E. coli* (taxid: 562)]

| Description | Scientific Name | Max Score | Total Score | Query Cover | E value | Per. ident | Acc. Len | Accession |
|---|---|---|---|---|---|---|---|---|
| dihydrolipoyllysine-residue succinyltransferase [Escherichia coli O157:H7] | Escherichia coli O157:H7 | 307 | 307 | 89% | 1.00E-103 | 67.08 | 240 | EFC3643281.1 |
| dihydrolipoyllysine-residue succinyltransferase [Escherichia coli] | Escherichia coli | 307 | 307 | 89% | 1.00E-103 | 67.08 | 235 | WP_250378311.1 |
| dihydrolipoyllysine-residue succinyltransferase [Escherichia coli] | Escherichia coli | 307 | 307 | 89% | 1.00E-103 | 67.08 | 234 | WP_149465242.1 |
| dihydrolipoyllysine-residue succinyltransferase [Escherichia coli O177] | Escherichia coli O177 | 307 | 307 | 89% | 2.00E-103 | 67.08 | 229 | EFA8854754.1 |
| dihydrolipoyllysine-residue succinyltransferase [Escherichia] | Escherichia | 307 | 307 | 89% | 2.00E-103 | 67.08 | 236 | WP_000629458.1 |
| TPA: dihydrolipoyllysine-residue succinyltransferase [Escherichia coli] | Escherichia coli | 307 | 307 | 89% | 2.00E-103 | 67.08 | 240 | HDY2775515.1 |
| dihydrolipoyllysine-residue succinyltransferase [Escherichia coli] | Escherichia coli | 306 | 306 | 89% | 2.00E-103 | 67.08 | 230 | EGE7843444.1 |
| dihydrolipoyllysine-residue succinyltransferase [Escherichia coli] | Escherichia coli | 307 | 307 | 89% | 2.00E-103 | 67.08 | 237 | WP_252389303.1 |
| dihydrolipoyllysine-residue succinyltransferase [Escherichia coli] | Escherichia coli | 307 | 307 | 89% | 2.00E-103 | 67.08 | 233 | WP_136823828.1 |
| dihydrolipoyllysine-residue succinyltransferase [Escherichia coli] | Escherichia coli | 307 | 307 | 89% | 2.00E-103 | 67.08 | 241 | EIH9592508.1 |
| dihydrolipoyllysine-residue succinyltransferase [Escherichia coli] | Escherichia coli | 307 | 307 | 89% | 2.00E-103 | 67.08 | 238 | EFO3983716.1 |
| dihydrolipoyllysine-residue succinyltransferase [Escherichia coli] | Escherichia coli | 307 | 307 | 89% | 2.00E-103 | 67.08 | 239 | EEZ3479170.1 |
| dihydrolipoyllysine-residue succinyltransferase [Escherichia coli O157:H7] | Escherichia coli O157:H7 | 306 | 306 | 89% | 2.00E-103 | 67.08 | 231 | EEW1174705.1 |
| dihydrolipoyllysine-residue succinyltransferase [Escherichia coli] | Escherichia coli | 306 | 306 | 89% | 2.00E-103 | 66.67 | 235 | WP_122792276.1 |
| dihydrolipoyllysine-residue succinyltransferase [Escherichia coli] | Escherichia coli | 307 | 307 | 89% | 2.00E-103 | 67.08 | 243 | TJB60735.1 |
| dihydrolipoyllysine-residue succinyltransferase [Escherichia coli] | Escherichia coli | 307 | 307 | 89% | 3.00E-103 | 67.08 | 248 | KAA0647049.1 |
| dihydrolipoyllysine-residue succinyltransferase [Escherichia coli] | Escherichia coli | 306 | 306 | 89% | 3.00E-103 | 67.36 | 228 | WP_229024141.1 |
| dihydrolipoyllysine-residue succinyltransferase [Escherichia coli] | Escherichia coli | 307 | 307 | 89% | 3.00E-103 | 67.08 | 257 | MCU7719730.1 |
| dihydrolipoyllysine-residue succinyltransferase [Escherichia coli] | Escherichia coli | 307 | 307 | 89% | 3.00E-103 | 67.08 | 259 | EIC1734575.1 |
| TPA: dihydrolipoyllysine-residue succinyltransferase [Escherichia coli] | Escherichia coli | 307 | 307 | 89% | 3.00E-103 | 67.08 | 261 | HAM4331889.1 |
| dihydrolipoyllysine-residue succinyltransferase [Escherichia coli] | Escherichia coli | 307 | 307 | 89% | 3.00E-103 | 67.08 | 255 | WP_042066964.1 |
| TPA: dihydrolipoyllysine-residue succinyltransferase [Escherichia coli] | Escherichia coli | 307 | 307 | 89% | 3.00E-103 | 67.08 | 249 | HBH7468286.1 |
| dihydrolipoyllysine-residue succinyltransferase [Escherichia coli] | Escherichia coli | 307 | 307 | 89% | 3.00E-103 | 67.08 | 250 | WP_323653019.1 |
| dihydrolipoyllysine-residue succinyltransferase [Escherichia coli] | Escherichia coli | 307 | 307 | 89% | 3.00E-103 | 67.08 | 256 | MCJ8642909.1 |

| | | | | | | | | |
|---|---|---|---|---|---|---|---|---|
| dihydrolipoyllysine-residue succinyltransferase [Escherichia coli] | Escherichia coli | 306 | 306 | 89% | 3.00E-103 | 67.08 | 242 | WP_149518815.1 |
| dihydrolipoyllysine-residue succinyltransferase [Escherichia coli] | Escherichia coli | 306 | 306 | 89% | 3.00E-103 | 67.08 | 243 | EEX5691720.1 |
| TPA: dihydrolipoyllysine-residue succinyltransferase [Escherichia coli] | Escherichia coli | 307 | 307 | 89% | 3.00E-103 | 67.08 | 254 | HDC0239447.1 |
| hypothetical protein ECZU26_29390 [Escherichia coli] | Escherichia coli | 307 | 307 | 89% | 3.00E-103 | 67.08 | 260 | GHL32114.1 |
| TPA: dihydrolipoyllysine-residue succinyltransferase [Escherichia coli] | Escherichia coli | 306 | 306 | 89% | 3.00E-103 | 67.08 | 244 | HBB1186217.1 |
| dihydrolipoyllysine-residue succinyltransferase [Escherichia coli] | Escherichia coli | 306 | 306 | 89% | 3.00E-103 | 67.08 | 245 | WP_136775487.1 |
| 2-oxoglutarate dehydrogenase complex dihydrolipoyllysine-residue succinyltransferase [Escherichia coli] | Escherichia coli | 308 | 308 | 89% | 4.00E-103 | 67.08 | 282 | MDY9040892.1 |
| dihydrolipoyllysine-residue succinyltransferase [Escherichia coli O145:H28] | Escherichia coli O145:H28 | 306 | 306 | 89% | 4.00E-103 | 67.08 | 246 | EJH5192981.1 |
| dihydrolipoyllysine-residue succinyltransferase [Escherichia coli] | Escherichia coli | 307 | 307 | 89% | 4.00E-103 | 67.08 | 260 | WP_252391012.1 |
| dihydrolipoyllysine-residue succinyltransferase [Escherichia coli] | Escherichia coli | 306 | 306 | 89% | 4.00E-103 | 67.08 | 247 | WP_185168965.1 |
| dihydrolipoyllysine-residue succinyltransferase [Escherichia coli] | Escherichia coli | 307 | 307 | 89% | 4.00E-103 | 67.08 | 263 | WP_047654718.1 |
| dihydrolipoamide succinyltransferase [Escherichia coli] | Escherichia coli | 307 | 307 | 89% | 4.00E-103 | 67.08 | 263 | OWF26202.1 |
| TPA: dihydrolipoyllysine-residue succinyltransferase [Escherichia coli] | Escherichia coli | 307 | 307 | 89% | 4.00E-103 | 67.08 | 264 | HAN7662089.1 |
| dihydrolipoyllysine-residue succinyltransferase [Escherichia coli] | Escherichia coli | 306 | 306 | 89% | 4.00E-103 | 67.08 | 248 | TJB30469.1 |
| dihydrolipoyllysine-residue succinyltransferase [Escherichia coli] | Escherichia coli | 307 | 307 | 89% | 4.00E-103 | 67.08 | 259 | EGE6907964.1 |
| dihydrolipoyllysine-residue succinyltransferase [Escherichia coli] | Escherichia coli | 307 | 307 | 89% | 4.00E-103 | 67.08 | 259 | WP_119682542.1 |
| dihydrolipoyllysine-residue succinyltransferase [Escherichia coli O145:H28] | Escherichia coli O145:H28 | 307 | 307 | 89% | 4.00E-103 | 67.08 | 264 | EJH5270977.1 |
| 2-oxoglutarate dehydrogenase complex dihydrolipoyllysine-residue succinyltransferase [Escherichia coli] | Escherichia coli | 309 | 309 | 89% | 4.00E-103 | 67.08 | 315 | WP_284108106.1 |
| 2-oxoglutarate dehydrogenase complex dihydrolipoyllysine-residue succinyltransferase [Escherichia coli] | Escherichia coli | 308 | 308 | 89% | 4.00E-103 | 67.08 | 280 | WP_320759799.1 |
| dihydrolipoyllysine-residue succinyltransferase [Escherichia coli] | Escherichia coli | 306 | 306 | 89% | 4.00E-103 | 67.08 | 252 | MDY9032137.1 |
| TPA: dihydrolipoyllysine-residue succinyltransferase [Escherichia coli] | Escherichia coli | 306 | 306 | 89% | 4.00E-103 | 67.08 | 252 | HBD1696035.1 |
| dihydrolipoyllysine-residue succinyltransferase [Escherichia coli] | Escherichia coli | 306 | 306 | 89% | 5.00E-103 | 67.08 | 253 | WP_136747018.1 |
| dihydrolipoyllysine-residue succinyltransferase [Escherichia coli] | Escherichia coli | 306 | 306 | 89% | 5.00E-103 | 67.08 | 253 | WP_136748022.1 |
| dihydrolipoyllysine-residue succinyltransferase [Escherichia coli] | Escherichia coli | 306 | 306 | 89% | 5.00E-103 | 67.08 | 253 | EFH6296858.1 |
| 2-oxoglutarate dehydrogenase complex dihydrolipoyllysine-residue succinyltransferase [Escherichia coli] | Escherichia coli | 308 | 308 | 89% | 5.00E-103 | 67.08 | 289 | WP_250216047.1 |
| dihydrolipoamide succinyltransferase component (E2) [Escherichia coli] | Escherichia coli | 307 | 307 | 89% | 5.00E-103 | 67.08 | 269 | VFT70387.1 |
| 2-oxoglutarate dehydrogenase complex dihydrolipoyllysine-residue succinyltransferase [Escherichia coli] | Escherichia coli | 307 | 307 | 89% | 5.00E-103 | 67.08 | 274 | MDY8964183.1 |

| | | | | | | | | |
|---|---|---|---|---|---|---|---|---|
| TPA: dihydrolipoyllysine-residue succinyltransferase [Escherichia coli] | Escherichia coli | 307 | 307 | 89% | 5.00E-103 | 67.08 | 265 | HAP0635211.1 |
| 2-oxoglutarate dehydrogenase complex dihydrolipoyllysine-residue succinyltransferase [Escherichia coli] | Escherichia coli | 307 | 307 | 89% | 5.00E-103 | 67.08 | 269 | WP_115969540.1 |
| 2-oxoglutarate dehydrogenase complex dihydrolipoyllysine-residue succinyltransferase [Escherichia coli] | Escherichia coli | 307 | 307 | 89% | 5.00E-103 | 67.08 | 269 | WP_249524430.1 |
| 2-oxoglutarate dehydrogenase complex dihydrolipoyllysine-residue succinyltransferase [Escherichia coli] | Escherichia coli | 308 | 308 | 89% | 5.00E-103 | 67.08 | 286 | WP_047081876.1 |
| 2-oxoglutarate dehydrogenase complex dihydrolipoyllysine-residue succinyltransferase [Escherichia coli] | Escherichia coli | 307 | 307 | 89% | 5.00E-103 | 67.08 | 274 | RIC38964.1 |
| dihydrolipoyllysine-residue succinyltransferase [Escherichia coli] | Escherichia coli | 306 | 306 | 89% | 5.00E-103 | 67.08 | 253 | WP_136801371.1 |
| 2-oxoglutarate dehydrogenase complex dihydrolipoyllysine-residue succinyltransferase [Escherichia coli] | Escherichia coli | 307 | 307 | 89% | 5.00E-103 | 67.08 | 270 | WP_250382326.1 |
| 2-oxoglutarate dehydrogenase complex dihydrolipoyllysine-residue succinyltransferase [Escherichia coli] | Escherichia coli | 308 | 308 | 89% | 6.00E-103 | 67.08 | 286 | WP_060643349.1 |
| 2-oxoglutarate dehydrogenase complex dihydrolipoyllysine-residue succinyltransferase [Escherichia coli] | Escherichia coli | 308 | 308 | 89% | 6.00E-103 | 67.08 | 295 | NAG00763.1 |
| 2-oxoglutarate dehydrogenase complex dihydrolipoyllysine-residue succinyltransferase [Escherichia coli] | Escherichia coli | 307 | 307 | 89% | 6.00E-103 | 67.08 | 275 | WP_136750337.1 |
| 2-oxoglutarate dehydrogenase complex dihydrolipoyllysine-residue succinyltransferase [Escherichia coli] | Escherichia coli | 308 | 308 | 89% | 6.00E-103 | 67.08 | 292 | WP_194498297.1 |
| dihydrolipoamide succinyltransferase component (E2) [Escherichia coli] | Escherichia coli | 307 | 307 | 89% | 6.00E-103 | 67.08 | 269 | STG69599.1 |
| 2-oxoglutarate dehydrogenase complex dihydrolipoyllysine-residue succinyltransferase [Escherichia coli] | Escherichia coli | 307 | 307 | 89% | 6.00E-103 | 67.08 | 287 | EEZ2060122.1 |
| TPA: 2-oxoglutarate dehydrogenase complex dihydrolipoyllysine-residue succinyltransferase [Escherichia coli] | Escherichia coli | 308 | 308 | 89% | 7.00E-103 | 67.08 | 312 | HCO0761733.1 |
| 2-oxoglutarate dehydrogenase complex dihydrolipoyllysine-residue succinyltransferase [Escherichia coli] | Escherichia coli | 308 | 308 | 89% | 8.00E-103 | 67.08 | 321 | EEY6693670.1 |
| 2-oxoglutarate dehydrogenase complex dihydrolipoyllysine-residue succinyltransferase [Escherichia coli] | Escherichia coli | 308 | 308 | 89% | 8.00E-103 | 67.08 | 324 | EER8331471.1 |
| 2-oxoglutarate dehydrogenase complex dihydrolipoyllysine-residue succinyltransferase [Escherichia coli] | Escherichia coli | 308 | 308 | 89% | 8.00E-103 | 67.08 | 324 | MDO4358214.1 |
| 2-oxoglutarate dehydrogenase complex dihydrolipoyllysine-residue succinyltransferase [Escherichia coli] | Escherichia coli | 308 | 308 | 89% | 8.00E-103 | 67.08 | 303 | EEY6784672.1 |
| 2-oxoglutarate dehydrogenase complex dihydrolipoyllysine-residue succinyltransferase [Escherichia coli] | Escherichia coli | 307 | 307 | 89% | 1.00E-102 | 67.08 | 300 | MBA1840094.1 |
| 2-oxoglutarate dehydrogenase complex dihydrolipoyllysine-residue succinyltransferase [Escherichia coli] | Escherichia coli | 308 | 308 | 89% | 1.00E-102 | 67.08 | 322 | WP_194154148.1 |
| 2-oxoglutarate dehydrogenase complex dihydrolipoyllysine-residue succinyltransferase [Escherichia coli] | Escherichia coli | 308 | 308 | 89% | 1.00E-102 | 67.08 | 322 | MCU8601182.1 |
| dihydrolipoamide succinyltransferase component (E2) [Escherichia coli] | Escherichia coli | 308 | 308 | 89% | 1.00E-102 | 67.08 | 332 | STM59242.1 |

| | | | | | | | | |
|---|---|---|---|---|---|---|---|---|
| 2-oxoglutarate dehydrogenase complex dihydrolipoyllysine-residue succinyltransferase [Escherichia coli] | Escherichia coli | 308 | 308 | 89% | 1.00E-102 | 67.08 | 318 | MCX0506393.1 |
| 2-oxoglutarate dehydrogenase complex dihydrolipoyllysine-residue succinyltransferase [Escherichia coli] | Escherichia coli | 308 | 308 | 89% | 1.00E-102 | 67.08 | 338 | WP_044071987.1 |
| 2-oxoglutarate dehydrogenase complex dihydrolipoyllysine-residue succinyltransferase [Escherichia coli] | Escherichia coli | 308 | 308 | 89% | 1.00E-102 | 67.08 | 337 | WP_048224949.1 |
| 2-oxoglutarate dehydrogenase complex dihydrolipoyllysine-residue succinyltransferase [Escherichia coli] | Escherichia coli | 308 | 308 | 89% | 1.00E-102 | 67.08 | 338 | MDY8712251.1 |
| 2-oxoglutarate dehydrogenase complex dihydrolipoyllysine-residue succinyltransferase [Escherichia coli] | Escherichia coli | 308 | 308 | 89% | 2.00E-102 | 67.08 | 328 | WP_229002757.1 |
| 2-oxoglutarate dehydrogenase complex dihydrolipoyllysine-residue succinyltransferase [Escherichia coli] | Escherichia coli | 311 | 311 | 89% | 2.00E-102 | 67.08 | 405 | UIU60256.1 |
| TPA: dihydrolipoyllysine-residue succinyltransferase [Escherichia coli] | Escherichia coli | 305 | 305 | 89% | 2.00E-102 | 67.08 | 250 | HBI3672065.1 |
| 2-oxoglutarate dehydrogenase complex dihydrolipoyllysine-residue succinyltransferase [Escherichia coli] | Escherichia coli | 308 | 308 | 89% | 2.00E-102 | 67.08 | 331 | WP_042068848.1 |
| TPA: 2-oxoglutarate dehydrogenase complex dihydrolipoyllysine-residue succinyltransferase [Escherichia coli] | Escherichia coli | 308 | 308 | 89% | 2.00E-102 | 67.08 | 332 | HCN8464961.1 |
| 2-oxoglutarate dehydrogenase complex dihydrolipoyllysine-residue succinyltransferase [Escherichia coli] | Escherichia coli | 310 | 310 | 89% | 2.00E-102 | 67.5 | 405 | EEZ9068234.1 |
| dihydrolipoyllysine-residue succinyltransferase [Escherichia coli] | Escherichia coli | 304 | 304 | 89% | 2.00E-102 | 67.23 | 227 | EIH2436553.1 |
| dihydrolipoyllysine-residue succinyltransferase [Escherichia coli] | Escherichia coli | 305 | 305 | 89% | 2.00E-102 | 67.08 | 253 | WP_347079825.1 |
| 2-oxoglutarate dehydrogenase complex dihydrolipoyllysine-residue succinyltransferase [Escherichia coli] | Escherichia coli | 310 | 310 | 89% | 2.00E-102 | 67.08 | 405 | ELJ3482170.1 |
| 2-oxoglutarate dehydrogenase complex dihydrolipoyllysine-residue succinyltransferase [Escherichia coli] | Escherichia coli | 310 | 310 | 89% | 2.00E-102 | 67.08 | 405 | WP_021515810.1 |
| 2-oxoglutarate dehydrogenase complex dihydrolipoyllysine-residue succinyltransferase [Escherichia coli] | Escherichia coli | 306 | 306 | 89% | 3.00E-102 | 67.08 | 279 | WP_247166683.1 |
| 2-oxoglutarate dehydrogenase complex dihydrolipoyllysine-residue succinyltransferase [Escherichia coli] | Escherichia coli | 308 | 308 | 89% | 3.00E-102 | 67.08 | 354 | MBE9780112.1 |
| 2-oxoglutarate dehydrogenase complex dihydrolipoyllysine-residue succinyltransferase [Escherichia coli] | Escherichia coli | 308 | 308 | 89% | 3.00E-102 | 67.08 | 345 | MBA1060735.1 |
| 2-oxoglutarate dehydrogenase complex dihydrolipoyllysine-residue succinyltransferase [Escherichia coli] | Escherichia coli | 310 | 310 | 89% | 4.00E-102 | 67.08 | 405 | WP_001704800.1 |
| 2-oxoglutarate dehydrogenase complex dihydrolipoyllysine-residue succinyltransferase [Escherichia coli] | Escherichia coli | 308 | 308 | 89% | 4.00E-102 | 67.08 | 366 | MXS49548.1 |
| 2-oxoglutarate dehydrogenase complex dihydrolipoyllysine-residue succinyltransferase [Escherichia coli] | Escherichia coli | 310 | 310 | 89% | 4.00E-102 | 67.08 | 405 | WP_033554859.1 |
| TPA: 2-oxoglutarate dehydrogenase complex dihydrolipoyllysine-residue succinyltransferase [Escherichia coli] | Escherichia coli | 310 | 310 | 89% | 4.00E-102 | 67.08 | 405 | HAX5241565.1 |
| 2-oxoglutarate dehydrogenase complex dihydrolipoyllysine-residue succinyltransferase [Escherichia coli] | Escherichia coli | 310 | 310 | 91% | 4.00E-102 | 66.39 | 405 | WP_136803128.1 |

| | | | | | | | | |
|---|---|---|---|---|---|---|---|---|
| 2-oxoglutarate dehydrogenase complex dihydrolipoyllysine-residue succinyltransferase [Escherichia coli] | Escherichia coli | 310 | 310 | 91% | 4.00E-102 | 66.39 | 405 | WP_096845185.1 |
| TPA: 2-oxoglutarate dehydrogenase complex dihydrolipoyllysine-residue succinyltransferase [Escherichia coli] | Escherichia coli | 310 | 310 | 89% | 4.00E-102 | 67.08 | 400 | HAX4889521.1 |
| 2-oxoglutarate dehydrogenase complex dihydrolipoyllysine-residue succinyltransferase [Escherichia coli] | Escherichia coli | 310 | 310 | 89% | 5.00E-102 | 67.5 | 405 | EFH5865371.1 |
| 2-oxoglutarate dehydrogenase complex dihydrolipoyllysine-residue succinyltransferase [Escherichia coli] | Escherichia coli | 310 | 310 | 89% | 5.00E-102 | 67.5 | 405 | EFL9867405.1 |
| 2-oxoglutarate dehydrogenase complex dihydrolipoyllysine-residue succinyltransferase [Escherichia coli] | Escherichia coli | 310 | 310 | 89% | 5.00E-102 | 67.5 | 405 | EIH5269730.1 |

Supplemental Table 6 | Protein BLAST results based on consensus sequence of automated model building on a 3.00 Å low-pass filtered volume map [Results limited to *E. coli* (taxid: 562)]

| Description | Scientific Name | Max Score | Total Score | Query Cover | E value | Per. ident | Acc. Len | Accession |
|---|---|---|---|---|---|---|---|---|
| dihydrolipoyllysine-residue succinyltransferase [Escherichia coli O157:H7] | Escherichia coli O157:H7 | 379 | 379 | 99% | 2.00E-132 | 81.62 | 231 | EEW1174705.1 |
| dihydrolipoyllysine-residue succinyltransferase [Escherichia coli] | Escherichia coli | 379 | 379 | 99% | 2.00E-132 | 81.62 | 239 | EEZ3479170.1 |
| dihydrolipoyllysine-residue succinyltransferase [Escherichia coli O157:H7] | Escherichia coli O157:H7 | 379 | 379 | 99% | 2.00E-132 | 81.62 | 240 | EFC3643281.1 |
| dihydrolipoyllysine-residue succinyltransferase [Escherichia coli] | Escherichia coli | 379 | 379 | 99% | 2.00E-132 | 81.62 | 235 | WP_250378311.1 |
| TPA: dihydrolipoyllysine-residue succinyltransferase [Escherichia coli] | Escherichia coli | 379 | 379 | 99% | 2.00E-132 | 81.62 | 240 | HDY2775515.1 |
| dihydrolipoyllysine-residue succinyltransferase [Escherichia] | Escherichia | 379 | 379 | 99% | 2.00E-132 | 81.62 | 236 | WP_000629458.1 |
| dihydrolipoyllysine-residue succinyltransferase [Escherichia coli] | Escherichia coli | 379 | 379 | 99% | 2.00E-132 | 81.62 | 235 | WP_122792276.1 |
| dihydrolipoyllysine-residue succinyltransferase [Escherichia coli] | Escherichia coli | 379 | 379 | 99% | 2.00E-132 | 81.62 | 233 | WP_136823828.1 |
| dihydrolipoyllysine-residue succinyltransferase [Escherichia coli] | Escherichia coli | 379 | 379 | 99% | 2.00E-132 | 81.62 | 234 | WP_149465242.1 |
| dihydrolipoyllysine-residue succinyltransferase [Escherichia coli] | Escherichia coli | 379 | 379 | 99% | 3.00E-132 | 81.62 | 241 | EIH9592508.1 |
| dihydrolipoyllysine-residue succinyltransferase [Escherichia coli] | Escherichia coli | 379 | 379 | 99% | 3.00E-132 | 81.62 | 243 | TJB60735.1 |
| dihydrolipoyllysine-residue succinyltransferase [Escherichia coli] | Escherichia coli | 379 | 379 | 99% | 3.00E-132 | 81.62 | 238 | EFO3983716.1 |
| dihydrolipoyllysine-residue succinyltransferase [Escherichia coli] | Escherichia coli | 379 | 379 | 99% | 3.00E-132 | 81.62 | 237 | WP_252389303.1 |
| dihydrolipoyllysine-residue succinyltransferase [Escherichia coli] | Escherichia coli | 379 | 379 | 99% | 3.00E-132 | 81.62 | 259 | EIC1734575.1 |
| dihydrolipoyllysine-residue succinyltransferase [Escherichia coli] | Escherichia coli | 379 | 379 | 99% | 4.00E-132 | 81.62 | 248 | KAA0647049.1 |
| TPA: dihydrolipoyllysine-residue succinyltransferase [Escherichia coli] | Escherichia coli | 379 | 379 | 99% | 4.00E-132 | 81.62 | 249 | HBH7468286.1 |
| dihydrolipoyllysine-residue succinyltransferase [Escherichia coli] | Escherichia coli | 379 | 379 | 99% | 4.00E-132 | 81.62 | 260 | WP_252391012.1 |
| 2-oxoglutarate dehydrogenase complex dihydrolipoyllysine-residue succinyltransferase [Escherichia coli] | Escherichia coli | 380 | 380 | 99% | 4.00E-132 | 81.62 | 282 | MDY9040892.1 |
| dihydrolipoyllysine-residue succinyltransferase [Escherichia coli] | Escherichia coli | 379 | 379 | 99% | 4.00E-132 | 81.62 | 259 | EGE6907964.1 |

| | | | | | | | | |
|---|---|---|---|---|---|---|---|---|
| dihydrolipoyllysine-residue succinyltransferase [Escherichia coli] | Escherichia coli | 379 | 379 | 99% | 4.00E-132 | 81.62 | 259 | WP_119682542.1 |
| hypothetical protein ECZU26_29390 [Escherichia coli] | Escherichia coli | 379 | 379 | 99% | 4.00E-132 | 81.62 | 260 | GHL32114.1 |
| TPA: dihydrolipoyllysine-residue succinyltransferase [Escherichia coli] | Escherichia coli | 379 | 379 | 99% | 4.00E-132 | 81.62 | 261 | HAM4331889.1 |
| dihydrolipoyllysine-residue succinyltransferase [Escherichia coli] | Escherichia coli | 379 | 379 | 99% | 4.00E-132 | 81.62 | 250 | WP_323653019.1 |
| dihydrolipoyllysine-residue succinyltransferase [Escherichia coli] | Escherichia coli | 379 | 379 | 99% | 5.00E-132 | 81.62 | 257 | MCU7719730.1 |
| dihydrolipoyllysine-residue succinyltransferase [Escherichia coli] | Escherichia coli | 379 | 379 | 99% | 5.00E-132 | 81.62 | 263 | WP_047654718.1 |
| dihydrolipoamide succinyltransferase [Escherichia coli] | Escherichia coli | 379 | 379 | 99% | 5.00E-132 | 81.62 | 263 | OWF26202.1 |
| dihydrolipoyllysine-residue succinyltransferase [Escherichia coli O145:H28] | Escherichia coli O145:H28 | 379 | 379 | 99% | 5.00E-132 | 81.62 | 264 | EJH5270977.1 |
| 2-oxoglutarate dehydrogenase complex dihydrolipoyllysine-residue succinyltransferase [Escherichia coli] | Escherichia coli | 380 | 380 | 99% | 5.00E-132 | 81.62 | 289 | WP_250216047.1 |
| dihydrolipoyllysine-residue succinyltransferase [Escherichia coli] | Escherichia coli | 378 | 378 | 99% | 5.00E-132 | 81.62 | 242 | WP_149518815.1 |
| TPA: dihydrolipoyllysine-residue succinyltransferase [Escherichia coli] | Escherichia coli | 379 | 379 | 99% | 5.00E-132 | 81.62 | 264 | HAN7662089.1 |
| dihydrolipoamide succinyltransferase component (E2) [Escherichia coli] | Escherichia coli | 379 | 379 | 99% | 5.00E-132 | 81.62 | 269 | VFT70387.1 |
| 2-oxoglutarate dehydrogenase complex dihydrolipoyllysine-residue succinyltransferase [Escherichia coli] | Escherichia coli | 384 | 384 | 99% | 5.00E-132 | 81.62 | 405 | UIU60256.1 |
| dihydrolipoyllysine-residue succinyltransferase [Escherichia coli] | Escherichia coli | 378 | 378 | 99% | 5.00E-132 | 81.62 | 243 | EEX5691720.1 |
| TPA: dihydrolipoyllysine-residue succinyltransferase [Escherichia coli] | Escherichia coli | 379 | 379 | 99% | 5.00E-132 | 81.62 | 254 | HDC0239447.1 |
| TPA: dihydrolipoyllysine-residue succinyltransferase [Escherichia coli] | Escherichia coli | 378 | 378 | 99% | 5.00E-132 | 81.62 | 244 | HBB1186217.1 |
| dihydrolipoyllysine-residue succinyltransferase [Escherichia coli] | Escherichia coli | 379 | 379 | 99% | 6.00E-132 | 81.62 | 255 | WP_042066964.1 |
| dihydrolipoyllysine-residue succinyltransferase [Escherichia coli] | Escherichia coli | 378 | 378 | 99% | 6.00E-132 | 81.62 | 245 | WP_136775487.1 |
| dihydrolipoyllysine-residue succinyltransferase [Escherichia coli O145:H28] | Escherichia coli O145:H28 | 378 | 378 | 99% | 6.00E-132 | 81.62 | 246 | EJH5192981.1 |
| dihydrolipoamide succinyltransferase component (E2) [Escherichia coli] | Escherichia coli | 379 | 379 | 99% | 6.00E-132 | 81.62 | 269 | STG69599.1 |
| dihydrolipoyllysine-residue succinyltransferase [Escherichia coli] | Escherichia coli | 378 | 378 | 99% | 6.00E-132 | 81.62 | 247 | WP_185168965.1 |
| TPA: dihydrolipoyllysine-residue succinyltransferase [Escherichia coli] | Escherichia coli | 379 | 379 | 99% | 6.00E-132 | 81.62 | 265 | HAP0635211.1 |
| dihydrolipoyllysine-residue succinyltransferase [Escherichia coli] | Escherichia coli | 379 | 379 | 99% | 6.00E-132 | 81.62 | 256 | MCJ8642909.1 |
| dihydrolipoyllysine-residue succinyltransferase [Escherichia coli] | Escherichia coli | 378 | 378 | 99% | 6.00E-132 | 81.62 | 248 | TJB30469.1 |
| 2-oxoglutarate dehydrogenase complex dihydrolipoyllysine-residue succinyltransferase [Escherichia coli] | Escherichia coli | 380 | 380 | 99% | 7.00E-132 | 81.62 | 286 | WP_060643349.1 |
| TPA: 2-oxoglutarate dehydrogenase complex dihydrolipoyllysine-residue succinyltransferase [Escherichia coli] | Escherichia coli | 384 | 384 | 99% | 7.00E-132 | 82.05 | 405 | HDV4762163.1 |
| 2-oxoglutarate dehydrogenase complex dihydrolipoyllysine-residue succinyltransferase [Escherichia coli] | Escherichia coli | 379 | 379 | 99% | 7.00E-132 | 81.62 | 286 | WP_047081876.1 |

| | | | | | | | | |
|---|---|---|---|---|---|---|---|---|
| 2-oxoglutarate dehydrogenase complex dihydrolipoyllysine-residue succinyltransferase [Escherichia coli] | Escherichia coli | 384 | 384 | 99% | 7.00E-132 | 82.05 | 405 | WP_103682200.1 |
| dihydrolipoyllysine-residue succinyltransferase [Escherichia coli] | Escherichia coli | 378 | 378 | 99% | 7.00E-132 | 81.62 | 252 | MDY9032137.1 |
| TPA: dihydrolipoyllysine-residue succinyltransferase [Escherichia coli] | Escherichia coli | 378 | 378 | 99% | 7.00E-132 | 81.62 | 252 | HBD1696035.1 |
| 2-oxoglutarate dehydrogenase complex dihydrolipoyllysine-residue succinyltransferase [Escherichia coli] | Escherichia coli | 379 | 379 | 99% | 7.00E-132 | 81.62 | 287 | EEZ2060122.1 |
| dihydrolipoyllysine-residue succinyltransferase [Escherichia coli] | Escherichia coli | 378 | 378 | 99% | 7.00E-132 | 81.62 | 253 | WP_136747018.1 |
| dihydrolipoyllysine-residue succinyltransferase [Escherichia coli] | Escherichia coli | 378 | 378 | 99% | 7.00E-132 | 81.62 | 253 | WP_136748022.1 |
| dihydrolipoyllysine-residue succinyltransferase [Escherichia coli] | Escherichia coli | 378 | 378 | 99% | 7.00E-132 | 81.62 | 253 | EFH6296858.1 |
| 2-oxoglutarate dehydrogenase complex dihydrolipoyllysine-residue succinyltransferase [Escherichia coli] | Escherichia coli | 381 | 381 | 99% | 8.00E-132 | 81.62 | 321 | EEY6693670.1 |
| 2-oxoglutarate dehydrogenase complex dihydrolipoyllysine-residue succinyltransferase [Escherichia coli] | Escherichia coli | 381 | 381 | 99% | 8.00E-132 | 81.62 | 324 | EER8331471.1 |
| 2-oxoglutarate dehydrogenase complex dihydrolipoyllysine-residue succinyltransferase [Escherichia coli] | Escherichia coli | 381 | 381 | 99% | 8.00E-132 | 81.62 | 324 | MDO4358214.1 |
| 2-oxoglutarate dehydrogenase complex dihydrolipoyllysine-residue succinyltransferase [Escherichia coli] | Escherichia coli | 379 | 379 | 99% | 8.00E-132 | 81.62 | 275 | WP_136750337.1 |
| 2-oxoglutarate dehydrogenase complex dihydrolipoyllysine-residue succinyltransferase [Escherichia coli] | Escherichia coli | 380 | 380 | 99% | 8.00E-132 | 81.62 | 303 | EEY6784672.1 |
| 2-oxoglutarate dehydrogenase complex dihydrolipoyllysine-residue succinyltransferase [Escherichia coli] | Escherichia coli | 384 | 384 | 99% | 8.00E-132 | 82.05 | 405 | EFD4584379.1 |
| 2-oxoglutarate dehydrogenase complex dihydrolipoyllysine-residue succinyltransferase [Escherichia coli] | Escherichia coli | 380 | 380 | 99% | 8.00E-132 | 81.62 | 295 | NAG00763.1 |
| 2-oxoglutarate dehydrogenase complex dihydrolipoyllysine-residue succinyltransferase [Escherichia coli] | Escherichia coli | 380 | 380 | 99% | 8.00E-132 | 81.62 | 322 | WP_194154148.1 |
| 2-oxoglutarate dehydrogenase complex dihydrolipoyllysine-residue succinyltransferase [Escherichia coli] | Escherichia coli | 380 | 380 | 99% | 8.00E-132 | 81.62 | 322 | MCU8601182.1 |
| TPA: 2-oxoglutarate dehydrogenase complex dihydrolipoyllysine-residue succinyltransferase [Escherichia coli] | Escherichia coli | 380 | 380 | 99% | 9.00E-132 | 81.62 | 312 | HCO0761733.1 |
| 2-oxoglutarate dehydrogenase complex dihydrolipoyllysine-residue succinyltransferase [Escherichia coli] | Escherichia coli | 380 | 380 | 99% | 9.00E-132 | 81.62 | 318 | MCX0506393.1 |
| dihydrolipoyllysine-residue succinyltransferase [Escherichia coli] | Escherichia coli | 378 | 378 | 99% | 9.00E-132 | 81.62 | 253 | WP_136801371.1 |
| 2-oxoglutarate dehydrogenase complex dihydrolipoyllysine-residue succinyltransferase [Escherichia coli] | Escherichia coli | 379 | 379 | 99% | 9.00E-132 | 81.62 | 270 | WP_250382326.1 |
| 2-oxoglutarate dehydrogenase complex dihydrolipoyllysine-residue succinyltransferase [Escherichia coli] | Escherichia coli | 379 | 379 | 99% | 9.00E-132 | 81.62 | 269 | WP_115969540.1 |
| 2-oxoglutarate dehydrogenase complex dihydrolipoyllysine-residue succinyltransferase [Escherichia coli] | Escherichia coli | 379 | 379 | 99% | 9.00E-132 | 81.62 | 269 | WP_249524430.1 |
| 2-oxoglutarate dehydrogenase complex dihydrolipoyllysine-residue succinyltransferase [Escherichia coli] | Escherichia coli | 379 | 379 | 99% | 9.00E-132 | 81.62 | 280 | WP_320759799.1 |

| | | | | | | | | |
|---|---|---|---|---|---|---|---|---|
| 2-oxoglutarate dehydrogenase complex dihydrolipoyllysine-residue succinyltransferase [Escherichia coli] | Escherichia coli | 379 | 379 | 99% | 1.00E-131 | 81.62 | 274 | RIC38964.1 |
| dihydrolipoamide succinyltransferase component (E2) [Escherichia coli] | Escherichia coli | 381 | 381 | 99% | 1.00E-131 | 81.62 | 332 | STM59242.1 |
| 2-oxoglutarate dehydrogenase complex dihydrolipoyllysine-residue succinyltransferase [Escherichia coli] | Escherichia coli | 380 | 380 | 99% | 1.00E-131 | 81.62 | 300 | MBA1840094.1 |
| 2-oxoglutarate dehydrogenase complex dihydrolipoyllysine-residue succinyltransferase [Escherichia coli] | Escherichia coli | 381 | 381 | 99% | 1.00E-131 | 81.62 | 338 | MDY8712251.1 |
| 2-oxoglutarate dehydrogenase complex dihydrolipoyllysine-residue succinyltransferase [Escherichia coli] | Escherichia coli | 384 | 384 | 99% | 1.00E-131 | 82.05 | 405 | EGX8818027.1 |
| 2-oxoglutarate dehydrogenase complex dihydrolipoyllysine-residue succinyltransferase [Escherichia coli] | Escherichia coli | 379 | 379 | 99% | 1.00E-131 | 81.62 | 274 | MDY8964183.1 |
| 2-oxoglutarate dehydrogenase complex dihydrolipoyllysine-residue succinyltransferase [Escherichia coli] | Escherichia coli | 379 | 379 | 99% | 1.00E-131 | 81.62 | 292 | WP_194498297.1 |
| 2-oxoglutarate dehydrogenase complex dihydrolipoyllysine-residue succinyltransferase [Escherichia coli] | Escherichia coli | 381 | 381 | 99% | 1.00E-131 | 81.62 | 337 | WP_048224949.1 |
| dihydrolipoyllysine-residue succinyltransferase [Escherichia coli] | Escherichia coli | 377 | 377 | 98% | 1.00E-131 | 81.55 | 230 | EGE7843444.1 |
| 2-oxoglutarate dehydrogenase complex dihydrolipoyllysine-residue succinyltransferase [Escherichia coli] | Escherichia coli | 383 | 383 | 99% | 1.00E-131 | 81.62 | 405 | EKM5464437.1 |
| 2-oxoglutarate dehydrogenase complex dihydrolipoyllysine-residue succinyltransferase [Escherichia coli] | Escherichia coli | 381 | 381 | 99% | 1.00E-131 | 81.62 | 338 | WP_044071987.1 |
| TPA: 2-oxoglutarate dehydrogenase complex dihydrolipoyllysine-residue succinyltransferase [Escherichia coli] | Escherichia coli | 383 | 383 | 99% | 1.00E-131 | 82.05 | 405 | HAN9520759.1 |
| 2-oxoglutarate dehydrogenase complex dihydrolipoyllysine-residue succinyltransferase [Escherichia coli] | Escherichia coli | 381 | 381 | 99% | 1.00E-131 | 81.62 | 354 | MBE9780112.1 |
| 2-oxoglutarate dehydrogenase complex dihydrolipoyllysine-residue succinyltransferase [Escherichia coli] | Escherichia coli | 380 | 380 | 99% | 2.00E-131 | 81.62 | 328 | WP_229002757.1 |
| TPA: 2-oxoglutarate dehydrogenase complex dihydrolipoyllysine-residue succinyltransferase [Escherichia coli] | Escherichia coli | 383 | 383 | 99% | 2.00E-131 | 81.62 | 400 | HDI9351051.1 |
| 2-oxoglutarate dehydrogenase complex dihydrolipoyllysine-residue succinyltransferase [Escherichia coli] | Escherichia coli | 383 | 383 | 99% | 2.00E-131 | 82.05 | 405 | WP_112916033.1 |
| 2-oxoglutarate dehydrogenase complex dihydrolipoyllysine-residue succinyltransferase [Escherichia coli] | Escherichia coli | 383 | 383 | 99% | 2.00E-131 | 82.05 | 405 | WP_024234840.1 |
| 2-oxoglutarate dehydrogenase complex dihydrolipoyllysine-residue succinyltransferase [Escherichia coli] | Escherichia coli | 383 | 383 | 99% | 2.00E-131 | 82.05 | 405 | WP_302422103.1 |
| TPA: 2-oxoglutarate dehydrogenase complex dihydrolipoyllysine-residue succinyltransferase [Escherichia coli] | Escherichia coli | 380 | 380 | 99% | 2.00E-131 | 81.62 | 332 | HCN8464961.1 |
| TPA: 2-oxoglutarate dehydrogenase complex dihydrolipoyllysine-residue succinyltransferase [Escherichia coli] | Escherichia coli | 383 | 383 | 99% | 2.00E-131 | 82.05 | 405 | HDS6732478.1 |
| dihydrolipoyllysine-residue succinyltransferase [Escherichia coli] | Escherichia coli | 383 | 383 | 99% | 2.00E-131 | 81.62 | 405 | EFO2789461.1 |
| TPA: 2-oxoglutarate dehydrogenase complex dihydrolipoyllysine-residue succinyltransferase [Escherichia coli] | Escherichia coli | 383 | 383 | 99% | 2.00E-131 | 81.62 | 405 | HEI3235152.1 |

| | | | | | | | | |
|---|---|---|---|---|---|---|---|---|
| 2-oxoglutarate dehydrogenase complex dihydrolipoyllysine-residue succinyltransferase [Escherichia coli] | Escherichia coli | 380 | 380 | 99% | 2.00E-131 | 81.62 | 331 | WP_042068848.1 |
| TPA: 2-oxoglutarate dehydrogenase complex dihydrolipoyllysine-residue succinyltransferase [Escherichia coli] | Escherichia coli | 383 | 383 | 99% | 2.00E-131 | 81.62 | 405 | HDK8920924.1 |
| TPA: 2-oxoglutarate dehydrogenase complex dihydrolipoyllysine-residue succinyltransferase [Escherichia coli] | Escherichia coli | 382 | 382 | 99% | 2.00E-131 | 81.62 | 375 | HBB3835421.1 |
| 2-oxoglutarate dehydrogenase complex dihydrolipoyllysine-residue succinyltransferase [Escherichia coli] | Escherichia coli | 381 | 381 | 99% | 2.00E-131 | 81.62 | 366 | MXS49548.1 |
| 2-oxoglutarate dehydrogenase complex dihydrolipoyllysine-residue succinyltransferase [Escherichia coli] | Escherichia coli | 383 | 383 | 99% | 2.00E-131 | 81.62 | 405 | WP_103572461.1 |
| 2-oxoglutarate dehydrogenase complex dihydrolipoyllysine-residue succinyltransferase [Escherichia coli] | Escherichia coli | 380 | 380 | 99% | 2.00E-131 | 81.62 | 345 | MBA1060735.1 |
| dihydrolipoamide succinyltransferase component (E2) [Escherichia coli] | Escherichia coli | 381 | 381 | 99% | 2.00E-131 | 81.62 | 370 | VED13043.1 |
| dihydrolipoyllysine-residue succinyltransferase, E2 component of oxoglutarate dehydrogenase (succinyl-transferring) complex [Escherichia coli 1827-70] | Escherichia coli 1827-70 | 381 | 381 | 99% | 2.00E-131 | 81.62 | 370 | EFQ01732.1 |
| 2-oxoglutarate dehydrogenase complex dihydrolipoyllysine-residue succinyltransferase [Escherichia coli] | Escherichia coli | 379 | 379 | 99% | 2.00E-131 | 81.62 | 315 | WP_284108106.1 |

### Supplemental Table 7 | Protein BLAST results based on consensus sequence of automated model building on a 4.00 Å low-pass filtered volume map [Results limited to *E. coli* (taxid: 562)]

| Description | Scientific Name | Max Score | Total Score | Query Cover | E value | Per. ident | Acc. Len | Accession |
|---|---|---|---|---|---|---|---|---|
| dihydrolipoyllysine-residue succinyltransferase [Escherichia coli] | Escherichia coli | 352 | 352 | 99% | 1.00E-121 | 73.62 | 239 | EEZ3479170.1 |
| dihydrolipoyllysine-residue succinyltransferase [Escherichia coli] | Escherichia coli | 352 | 352 | 99% | 1.00E-121 | 73.62 | 235 | WP_250378311.1 |
| dihydrolipoyllysine-residue succinyltransferase [Escherichia coli] | Escherichia coli | 352 | 352 | 99% | 1.00E-121 | 73.62 | 238 | EFO3983716.1 |
| dihydrolipoyllysine-residue succinyltransferase [Escherichia] | Escherichia | 352 | 352 | 99% | 1.00E-121 | 73.62 | 236 | WP_000629458.1 |
| TPA: dihydrolipoyllysine-residue succinyltransferase [Escherichia coli] | Escherichia coli | 352 | 352 | 99% | 2.00E-121 | 73.62 | 240 | HDY2775515.1 |
| dihydrolipoyllysine-residue succinyltransferase [Escherichia coli O157:H7] | Escherichia coli O157:H7 | 352 | 352 | 99% | 2.00E-121 | 73.62 | 240 | EFC3643281.1 |
| dihydrolipoyllysine-residue succinyltransferase [Escherichia coli] | Escherichia coli | 351 | 351 | 99% | 2.00E-121 | 73.62 | 234 | WP_149465242.1 |
| dihydrolipoyllysine-residue succinyltransferase [Escherichia coli] | Escherichia coli | 351 | 351 | 99% | 2.00E-121 | 73.62 | 233 | WP_136823828.1 |
| dihydrolipoyllysine-residue succinyltransferase [Escherichia coli] | Escherichia coli | 351 | 351 | 99% | 2.00E-121 | 73.62 | 237 | WP_252389303.1 |
| dihydrolipoyllysine-residue succinyltransferase [Escherichia coli] | Escherichia coli | 351 | 351 | 99% | 2.00E-121 | 73.62 | 241 | EIH9592508.1 |
| dihydrolipoyllysine-residue succinyltransferase [Escherichia coli] | Escherichia coli | 352 | 352 | 99% | 2.00E-121 | 73.62 | 243 | TJB60735.1 |
| dihydrolipoyllysine-residue succinyltransferase [Escherichia coli] | Escherichia coli | 351 | 351 | 99% | 2.00E-121 | 73.19 | 235 | WP_122792276.1 |

| | | | | | | | | |
|---|---|---|---|---|---|---|---|---|
| dihydrolipoyllysine-residue succinyltransferase [Escherichia coli] | Escherichia coli | 351 | 351 | 98% | 3.00E-121 | 73.93 | 242 | WP_149518815.1 |
| dihydrolipoyllysine-residue succinyltransferase [Escherichia coli] | Escherichia coli | 351 | 351 | 98% | 3.00E-121 | 73.93 | 243 | EEX5691720.1 |
| dihydrolipoyllysine-residue succinyltransferase [Escherichia coli] | Escherichia coli | 351 | 351 | 98% | 3.00E-121 | 73.93 | 248 | KAA0647049.1 |
| TPA: dihydrolipoyllysine-residue succinyltransferase [Escherichia coli] | Escherichia coli | 351 | 351 | 98% | 3.00E-121 | 73.93 | 244 | HBB1186217.1 |
| dihydrolipoyllysine-residue succinyltransferase [Escherichia coli] | Escherichia coli | 352 | 352 | 98% | 3.00E-121 | 73.93 | 257 | MCU7719730.1 |
| dihydrolipoyllysine-residue succinyltransferase [Escherichia coli] | Escherichia coli | 351 | 351 | 98% | 3.00E-121 | 73.93 | 245 | WP_136775487.1 |
| TPA: dihydrolipoyllysine-residue succinyltransferase [Escherichia coli] | Escherichia coli | 351 | 351 | 98% | 3.00E-121 | 73.93 | 249 | HBH7468286.1 |
| dihydrolipoyllysine-residue succinyltransferase [Escherichia coli O145:H28] | Escherichia coli O145:H28 | 351 | 351 | 98% | 3.00E-121 | 73.93 | 246 | EJH5192981.1 |
| dihydrolipoyllysine-residue succinyltransferase [Escherichia coli] | Escherichia coli | 351 | 351 | 98% | 4.00E-121 | 73.93 | 247 | WP_185168965.1 |
| dihydrolipoyllysine-residue succinyltransferase [Escherichia coli] | Escherichia coli | 351 | 351 | 98% | 4.00E-121 | 73.93 | 259 | EGE6907964.1 |
| dihydrolipoyllysine-residue succinyltransferase [Escherichia coli] | Escherichia coli | 351 | 351 | 98% | 4.00E-121 | 73.93 | 259 | WP_119682542.1 |
| dihydrolipoyllysine-residue succinyltransferase [Escherichia coli] | Escherichia coli | 351 | 351 | 98% | 4.00E-121 | 73.93 | 248 | TJB30469.1 |
| dihydrolipoyllysine-residue succinyltransferase [Escherichia coli] | Escherichia coli | 351 | 351 | 98% | 4.00E-121 | 73.93 | 259 | EIC1734575.1 |
| dihydrolipoyllysine-residue succinyltransferase [Escherichia coli] | Escherichia coli | 351 | 351 | 98% | 4.00E-121 | 73.93 | 260 | WP_252391012.1 |
| TPA: dihydrolipoyllysine-residue succinyltransferase [Escherichia coli] | Escherichia coli | 351 | 351 | 98% | 4.00E-121 | 73.93 | 254 | HDC0239447.1 |
| dihydrolipoyllysine-residue succinyltransferase [Escherichia coli] | Escherichia coli | 351 | 351 | 98% | 4.00E-121 | 73.93 | 255 | WP_042066964.1 |
| dihydrolipoyllysine-residue succinyltransferase [Escherichia coli O145:H28] | Escherichia coli O145:H28 | 352 | 352 | 98% | 4.00E-121 | 73.93 | 264 | EJH5270977.1 |
| hypothetical protein ECZU26_29390 [Escherichia coli] | Escherichia coli | 351 | 351 | 98% | 4.00E-121 | 73.93 | 260 | GHL32114.1 |
| dihydrolipoyllysine-residue succinyltransferase [Escherichia coli] | Escherichia coli | 351 | 351 | 98% | 4.00E-121 | 73.93 | 250 | WP_323653019.1 |
| dihydrolipoamide succinyltransferase component (E2) [Escherichia coli] | Escherichia coli | 352 | 352 | 98% | 4.00E-121 | 73.93 | 269 | VFT70387.1 |
| dihydrolipoyllysine-residue succinyltransferase [Escherichia coli] | Escherichia coli | 351 | 351 | 98% | 4.00E-121 | 73.93 | 252 | MDY9032137.1 |
| TPA: dihydrolipoyllysine-residue succinyltransferase [Escherichia coli] | Escherichia coli | 351 | 351 | 98% | 4.00E-121 | 73.93 | 252 | HBD1696035.1 |
| dihydrolipoamide succinyltransferase component (E2) [Escherichia coli] | Escherichia coli | 352 | 352 | 98% | 4.00E-121 | 73.93 | 269 | STG69599.1 |
| dihydrolipoyllysine-residue succinyltransferase [Escherichia coli] | Escherichia coli | 351 | 351 | 98% | 4.00E-121 | 73.93 | 253 | WP_136747018.1 |
| dihydrolipoyllysine-residue succinyltransferase [Escherichia coli] | Escherichia coli | 351 | 351 | 98% | 4.00E-121 | 73.93 | 253 | WP_136748022.1 |
| dihydrolipoyllysine-residue succinyltransferase [Escherichia coli] | Escherichia coli | 351 | 351 | 98% | 4.00E-121 | 73.93 | 253 | EFH6296858.1 |

| | | | | | | | | |
|---|---|---|---|---|---|---|---|---|
| TPA: dihydrolipoyllysine-residue succinyltransferase [Escherichia coli] | Escherichia coli | 351 | 351 | 98% | 4.00E-121 | 73.93 | 261 | HAM4331889.1 |
| TPA: dihydrolipoyllysine-residue succinyltransferase [Escherichia coli] | Escherichia coli | 351 | 351 | 98% | 5.00E-121 | 73.93 | 265 | HAP0635211.1 |
| dihydrolipoyllysine-residue succinyltransferase [Escherichia coli] | Escherichia coli | 351 | 351 | 98% | 5.00E-121 | 73.93 | 256 | MCJ8642909.1 |
| 2-oxoglutarate dehydrogenase complex dihydrolipoyllysine-residue succinyltransferase [Escherichia coli] | Escherichia coli | 352 | 352 | 98% | 5.00E-121 | 73.93 | 269 | WP_115969540.1 |
| 2-oxoglutarate dehydrogenase complex dihydrolipoyllysine-residue succinyltransferase [Escherichia coli] | Escherichia coli | 352 | 352 | 98% | 5.00E-121 | 73.93 | 269 | WP_249524430.1 |
| 2-oxoglutarate dehydrogenase complex dihydrolipoyllysine-residue succinyltransferase [Escherichia coli] | Escherichia coli | 352 | 352 | 98% | 5.00E-121 | 73.93 | 282 | MDY9040892.1 |
| 2-oxoglutarate dehydrogenase complex dihydrolipoyllysine-residue succinyltransferase [Escherichia coli] | Escherichia coli | 351 | 351 | 98% | 6.00E-121 | 73.93 | 270 | WP_250382326.1 |
| 2-oxoglutarate dehydrogenase complex dihydrolipoyllysine-residue succinyltransferase [Escherichia coli] | Escherichia coli | 352 | 352 | 98% | 6.00E-121 | 73.93 | 280 | WP_320759799.1 |
| dihydrolipoyllysine-residue succinyltransferase [Escherichia coli] | Escherichia coli | 351 | 351 | 98% | 6.00E-121 | 73.93 | 263 | WP_047654718.1 |
| dihydrolipoamide succinyltransferase [Escherichia coli] | Escherichia coli | 351 | 351 | 98% | 6.00E-121 | 73.93 | 263 | OWF26202.1 |
| TPA: dihydrolipoyllysine-residue succinyltransferase [Escherichia coli] | Escherichia coli | 351 | 351 | 98% | 6.00E-121 | 73.93 | 264 | HAN7662089.1 |
| 2-oxoglutarate dehydrogenase complex dihydrolipoyllysine-residue succinyltransferase [Escherichia coli] | Escherichia coli | 351 | 351 | 98% | 6.00E-121 | 73.93 | 274 | MDY8964183.1 |
| 2-oxoglutarate dehydrogenase complex dihydrolipoyllysine-residue succinyltransferase [Escherichia coli] | Escherichia coli | 352 | 352 | 98% | 6.00E-121 | 73.93 | 289 | WP_250216047.1 |
| 2-oxoglutarate dehydrogenase complex dihydrolipoyllysine-residue succinyltransferase [Escherichia coli] | Escherichia coli | 352 | 352 | 98% | 7.00E-121 | 73.93 | 287 | EEZ2060122.1 |
| 2-oxoglutarate dehydrogenase complex dihydrolipoyllysine-residue succinyltransferase [Escherichia coli] | Escherichia coli | 356 | 356 | 99% | 7.00E-121 | 74.04 | 405 | WP_136803128.1 |
| 2-oxoglutarate dehydrogenase complex dihydrolipoyllysine-residue succinyltransferase [Escherichia coli] | Escherichia coli | 356 | 356 | 99% | 7.00E-121 | 74.04 | 405 | WP_096845185.1 |
| dihydrolipoyllysine-residue succinyltransferase [Escherichia coli] | Escherichia coli | 350 | 350 | 98% | 7.00E-121 | 73.93 | 253 | WP_136801371.1 |
| 2-oxoglutarate dehydrogenase complex dihydrolipoyllysine-residue succinyltransferase [Escherichia coli] | Escherichia coli | 352 | 352 | 98% | 7.00E-121 | 73.93 | 286 | WP_060643349.1 |
| 2-oxoglutarate dehydrogenase complex dihydrolipoyllysine-residue succinyltransferase [Escherichia coli] | Escherichia coli | 351 | 351 | 98% | 8.00E-121 | 73.93 | 274 | RIC38964.1 |
| 2-oxoglutarate dehydrogenase complex dihydrolipoyllysine-residue succinyltransferase [Escherichia coli] | Escherichia coli | 351 | 351 | 98% | 8.00E-121 | 73.93 | 275 | WP_136750337.1 |
| TPA: 2-oxoglutarate dehydrogenase complex dihydrolipoyllysine-residue succinyltransferase [Escherichia coli] | Escherichia coli | 352 | 352 | 98% | 9.00E-121 | 73.93 | 312 | HCO0761733.1 |

| | | | | | | | | |
|---|---|---|---|---|---|---|---|---|
| 2-oxoglutarate dehydrogenase complex dihydrolipoyllysine-residue succinyltransferase [Escherichia coli] | Escherichia coli | 352 | 352 | 98% | 9.00E-121 | 73.93 | 295 | NAG00763.1 |
| 2-oxoglutarate dehydrogenase complex dihydrolipoyllysine-residue succinyltransferase [Escherichia coli] | Escherichia coli | 356 | 356 | 99% | 9.00E-121 | 74.04 | 405 | MDY9786167.1 |
| 2-oxoglutarate dehydrogenase complex dihydrolipoyllysine-residue succinyltransferase [Escherichia coli] | Escherichia coli | 352 | 352 | 98% | 9.00E-121 | 73.93 | 292 | WP_194498297.1 |
| 2-oxoglutarate dehydrogenase complex dihydrolipoyllysine-residue succinyltransferase [Escherichia coli] | Escherichia coli | 352 | 352 | 98% | 9.00E-121 | 73.93 | 303 | EEY6784672.1 |
| 2-oxoglutarate dehydrogenase complex dihydrolipoyllysine-residue succinyltransferase [Escherichia coli] | Escherichia coli | 351 | 351 | 98% | 9.00E-121 | 73.93 | 286 | WP_047081876.1 |
| 2-oxoglutarate dehydrogenase complex dihydrolipoyllysine-residue succinyltransferase [Escherichia coli] | Escherichia coli | 353 | 353 | 98% | 1.00E-120 | 73.93 | 321 | EEY6693670.1 |
| 2-oxoglutarate dehydrogenase complex dihydrolipoyllysine-residue succinyltransferase [Escherichia coli] | Escherichia coli | 356 | 356 | 98% | 1.00E-120 | 73.93 | 405 | WP_192459758.1 |
| 2-oxoglutarate dehydrogenase complex dihydrolipoyllysine-residue succinyltransferase [Escherichia coli] | Escherichia coli | 353 | 353 | 98% | 1.00E-120 | 73.93 | 324 | EER8331471.1 |
| 2-oxoglutarate dehydrogenase complex dihydrolipoyllysine-residue succinyltransferase [Escherichia coli] | Escherichia coli | 353 | 353 | 98% | 1.00E-120 | 73.93 | 324 | MDO4358214.1 |
| TPA: dihydrolipoyllysine-residue succinyltransferase [Escherichia coli] | Escherichia coli | 350 | 350 | 99% | 1.00E-120 | 73.62 | 250 | HBI3672065.1 |
| 2-oxoglutarate dehydrogenase complex dihydrolipoyllysine-residue succinyltransferase [Escherichia coli] | Escherichia coli | 352 | 352 | 98% | 1.00E-120 | 73.93 | 300 | MBA1840094.1 |
| 2-oxoglutarate dehydrogenase complex dihydrolipoyllysine-residue succinyltransferase [Escherichia coli] | Escherichia coli | 353 | 353 | 98% | 1.00E-120 | 73.93 | 338 | MDY8712251.1 |
| dihydrolipoyllysine-residue succinyltransferase [Escherichia coli O157:H7] | Escherichia coli O157:H7 | 349 | 349 | 98% | 1.00E-120 | 73.82 | 231 | EEW1174705.1 |
| 2-oxoglutarate dehydrogenase complex dihydrolipoyllysine-residue succinyltransferase [Escherichia coli] | Escherichia coli | 352 | 352 | 98% | 1.00E-120 | 73.93 | 318 | MCX0506393.1 |
| dihydrolipoamide succinyltransferase component (E2) [Escherichia coli] | Escherichia coli | 353 | 353 | 98% | 1.00E-120 | 73.93 | 332 | STM59242.1 |
| 2-oxoglutarate dehydrogenase complex dihydrolipoyllysine-residue succinyltransferase [Escherichia coli] | Escherichia coli | 352 | 352 | 98% | 1.00E-120 | 73.93 | 322 | WP_194154148.1 |
| 2-oxoglutarate dehydrogenase complex dihydrolipoyllysine-residue succinyltransferase [Escherichia coli] | Escherichia coli | 352 | 352 | 98% | 1.00E-120 | 73.93 | 322 | MCU8601182.1 |
| 2-oxoglutarate dehydrogenase complex dihydrolipoyllysine-residue succinyltransferase [Escherichia coli] | Escherichia coli | 353 | 353 | 98% | 1.00E-120 | 73.93 | 328 | WP_229002757.1 |
| 2-oxoglutarate dehydrogenase complex dihydrolipoyllysine-residue succinyltransferase [Escherichia coli] | Escherichia coli | 355 | 355 | 98% | 1.00E-120 | 74.36 | 405 | MDF1405070.1 |
| 2-oxoglutarate dehydrogenase complex dihydrolipoyllysine-residue succinyltransferase [Escherichia coli] | Escherichia coli | 355 | 355 | 98% | 1.00E-120 | 74.36 | 405 | EEZ9068234.1 |

| | | | | | | | | |
|---|---|---|---|---|---|---|---|---|
| TPA: 2-oxoglutarate dehydrogenase complex dihydrolipoyllysine-residue succinyltransferase [Escherichia coli] | Escherichia coli | 353 | 353 | 98% | 2.00E-120 | 73.93 | 332 | HCN8464961.1 |
| 2-oxoglutarate dehydrogenase complex dihydrolipoyllysine-residue succinyltransferase [Escherichia coli] | Escherichia coli | 353 | 353 | 98% | 2.00E-120 | 73.93 | 337 | WP_048224949.1 |
| 2-oxoglutarate dehydrogenase complex dihydrolipoyllysine-residue succinyltransferase [Escherichia coli] | Escherichia coli | 352 | 352 | 98% | 2.00E-120 | 73.93 | 331 | WP_042068848.1 |
| 2-oxoglutarate dehydrogenase complex dihydrolipoyllysine-residue succinyltransferase [Escherichia coli] | Escherichia coli | 353 | 353 | 98% | 2.00E-120 | 73.93 | 345 | MBA1060735.1 |
| 2-oxoglutarate dehydrogenase complex dihydrolipoyllysine-residue succinyltransferase [Escherichia coli] | Escherichia coli | 353 | 353 | 98% | 2.00E-120 | 73.93 | 354 | MBE9780112.1 |
| 2-oxoglutarate dehydrogenase complex dihydrolipoyllysine-residue succinyltransferase [Escherichia coli] | Escherichia coli | 355 | 355 | 98% | 2.00E-120 | 73.93 | 405 | EKM5464437.1 |
| 2-oxoglutarate dehydrogenase complex dihydrolipoyllysine-residue succinyltransferase [Escherichia coli] | Escherichia coli | 353 | 353 | 98% | 2.00E-120 | 73.93 | 338 | WP_044071987.1 |
| 2-oxoglutarate dehydrogenase complex dihydrolipoyllysine-residue succinyltransferase [Escherichia coli] | Escherichia coli | 355 | 355 | 98% | 2.00E-120 | 74.36 | 405 | WP_032210407.1 |
| TPA: 2-oxoglutarate dehydrogenase complex dihydrolipoyllysine-residue succinyltransferase [Escherichia coli] | Escherichia coli | 355 | 355 | 98% | 2.00E-120 | 74.36 | 405 | HAH8524627.1 |
| dihydrolipoyllysine-residue succinyltransferase [Escherichia coli] | Escherichia coli | 349 | 349 | 98% | 2.00E-120 | 73.93 | 253 | WP_347079825.1 |
| 2-oxoglutarate dehydrogenase complex dihydrolipoyllysine-residue succinyltransferase [Escherichia coli] | Escherichia coli | 355 | 355 | 98% | 2.00E-120 | 74.36 | 405 | EIR6544570.1 |
| TPA: 2-oxoglutarate dehydrogenase complex dihydrolipoyllysine-residue succinyltransferase [Escherichia coli] | Escherichia coli | 355 | 355 | 98% | 2.00E-120 | 73.93 | 405 | HCP8029924.1 |
| 2-oxoglutarate dehydrogenase complex dihydrolipoyllysine-residue succinyltransferase [Escherichia] | Escherichia | 355 | 355 | 98% | 2.00E-120 | 74.36 | 405 | WP_000099817.1 |
| 2-oxoglutarate dehydrogenase complex dihydrolipoyllysine-residue succinyltransferase [Escherichia coli] | Escherichia coli | 350 | 350 | 98% | 2.00E-120 | 73.93 | 279 | WP_247166683.1 |
| 2-oxoglutarate dehydrogenase complex dihydrolipoyllysine-residue succinyltransferase [Escherichia coli] | Escherichia coli | 355 | 355 | 99% | 2.00E-120 | 73.31 | 405 | UIU60256.1 |
| TPA: 2-oxoglutarate dehydrogenase complex dihydrolipoyllysine-residue succinyltransferase [Escherichia coli] | Escherichia coli | 355 | 355 | 98% | 2.00E-120 | 74.36 | 405 | HCT2454644.1 |
| 2-oxoglutarate dehydrogenase complex dihydrolipoyllysine-residue succinyltransferase [Escherichia coli] | Escherichia coli | 353 | 353 | 98% | 2.00E-120 | 73.93 | 366 | MXS49548.1 |
| 2-oxoglutarate dehydrogenase complex dihydrolipoyllysine-residue succinyltransferase [Escherichia coli] | Escherichia coli | 355 | 355 | 98% | 2.00E-120 | 74.36 | 405 | WP_112916033.1 |
| TPA: 2-oxoglutarate dehydrogenase complex dihydrolipoyllysine-residue succinyltransferase [Escherichia coli] | Escherichia coli | 355 | 355 | 98% | 2.00E-120 | 74.36 | 405 | HAN9520759.1 |

| | | | | | | | | |
|---|---|---|---|---|---|---|---|---|
| 2-oxoglutarate dehydrogenase complex dihydrolipoyllysine-residue succinyltransferase [Escherichia coli] | Escherichia coli | 355 | 355 | 98% | 2.00E-120 | 74.36 | 405 | WP_024234840.1 |
| 2-oxoglutarate dehydrogenase complex dihydrolipoyllysine-residue succinyltransferase [Escherichia coli] | Escherichia coli | 355 | 355 | 98% | 2.00E-120 | 74.36 | 405 | WP_302422103.1 |

### Supplemental Table 8 | Protein BLAST results based on consensus sequence of automated model building on a 4.50 Å low-pass filtered volume map [Results limited to *E. coli* (taxid: 562)]

| Description | Scientific Name | Max Score | Total Score | Query Cover | E value | Per. ident | Acc. Len | Accession |
|---|---|---|---|---|---|---|---|---|
| dihydrolipoyllysine-residue succinyltransferase [Escherichia coli] | Escherichia coli | 132 | 132 | 97% | 5.00E-36 | 46.15 | 218 | MCJ8727037.1 |
| dihydrolipoyllysine-residue succinyltransferase [Escherichia coli] | Escherichia coli | 132 | 132 | 97% | 8.00E-36 | 46.15 | 234 | WP_149465242.1 |
| dihydrolipoyllysine-residue succinyltransferase [Escherichia coli] | Escherichia coli | 131 | 131 | 97% | 8.00E-36 | 46.15 | 233 | WP_136823828.1 |
| dihydrolipoyllysine-residue succinyltransferase [Escherichia coli] | Escherichia coli | 131 | 131 | 97% | 8.00E-36 | 46.15 | 235 | WP_250378311.1 |
| dihydrolipoyllysine-residue succinyltransferase [Escherichia] | Escherichia | 131 | 131 | 97% | 9.00E-36 | 46.15 | 236 | WP_000629458.1 |
| dihydrolipoyllysine-residue succinyltransferase [Escherichia coli O157:H7] | Escherichia coli O157:H7 | 131 | 131 | 97% | 9.00E-36 | 46.15 | 231 | EEW1174705.1 |
| dihydrolipoyllysine-residue succinyltransferase [Escherichia coli] | Escherichia coli | 131 | 131 | 97% | 9.00E-36 | 46.15 | 237 | WP_252389303.1 |
| TPA: dihydrolipoyllysine-residue succinyltransferase [Escherichia coli] | Escherichia coli | 131 | 131 | 97% | 1.00E-35 | 46.15 | 240 | HDY2775515.1 |
| dihydrolipoyllysine-residue succinyltransferase [Escherichia coli O157:H7] | Escherichia coli O157:H7 | 131 | 131 | 97% | 1.00E-35 | 46.15 | 240 | EFC3643281.1 |
| dihydrolipoyllysine-residue succinyltransferase [Escherichia coli] | Escherichia coli | 131 | 131 | 97% | 1.00E-35 | 46.15 | 241 | EIH9592508.1 |
| dihydrolipoyllysine-residue succinyltransferase [Escherichia coli] | Escherichia coli | 131 | 131 | 97% | 1.00E-35 | 46.15 | 238 | EFO3983716.1 |
| dihydrolipoyllysine-residue succinyltransferase [Escherichia coli] | Escherichia coli | 131 | 131 | 97% | 1.00E-35 | 46.15 | 239 | EEZ3479170.1 |
| dihydrolipoyllysine-residue succinyltransferase [Escherichia coli] | Escherichia coli | 131 | 131 | 97% | 1.00E-35 | 46.15 | 235 | WP_122792276.1 |
| dihydrolipoyllysine-residue succinyltransferase [Escherichia coli] | Escherichia coli | 132 | 132 | 97% | 1.00E-35 | 46.15 | 248 | KAA0647049.1 |
| dihydrolipoyllysine-residue succinyltransferase [Escherichia coli] | Escherichia coli | 131 | 131 | 97% | 1.00E-35 | 46.15 | 243 | TJB60735.1 |
| dihydrolipoyllysine-residue succinyltransferase [Escherichia coli] | Escherichia coli | 131 | 131 | 97% | 1.00E-35 | 46.15 | 242 | WP_149518815.1 |
| dihydrolipoyllysine-residue succinyltransferase [Escherichia coli] | Escherichia coli | 131 | 131 | 97% | 1.00E-35 | 46.15 | 243 | EEX5691720.1 |
| TPA: dihydrolipoyllysine-residue succinyltransferase [Escherichia coli] | Escherichia coli | 131 | 131 | 97% | 1.00E-35 | 46.15 | 244 | HBB1186217.1 |
| dihydrolipoyllysine-residue succinyltransferase [Escherichia coli] | Escherichia coli | 131 | 131 | 97% | 1.00E-35 | 46.15 | 245 | WP_136775487.1 |
| dihydrolipoyllysine-residue succinyltransferase [Escherichia coli O145:H28] | Escherichia coli O145:H28 | 131 | 131 | 97% | 1.00E-35 | 46.15 | 246 | EJH5192981.1 |

| | | | | | | | | |
|---|---|---|---|---|---|---|---|---|
| dihydrolipoyllysine-residue succinyltransferase [Escherichia coli] | Escherichia coli | 131 | 131 | 97% | 1.00E-35 | 46.15 | 247 | WP_185168965.1 |
| dihydrolipoyllysine-residue succinyltransferase [Escherichia coli] | Escherichia coli | 131 | 131 | 97% | 1.00E-35 | 46.15 | 248 | TJB30469.1 |
| dihydrolipoyllysine-residue succinyltransferase [Escherichia coli] | Escherichia coli | 132 | 132 | 97% | 1.00E-35 | 46.15 | 256 | MCJ8642909.1 |
| TPA: dihydrolipoyllysine-residue succinyltransferase [Escherichia coli] | Escherichia coli | 131 | 131 | 97% | 1.00E-35 | 46.15 | 249 | HBH7468286.1 |
| dihydrolipoyllysine-residue succinyltransferase [Escherichia coli] | Escherichia coli | 131 | 131 | 97% | 1.00E-35 | 46.15 | 250 | WP_323653019.1 |
| dihydrolipoyllysine-residue succinyltransferase [Escherichia coli] | Escherichia coli | 131 | 131 | 97% | 1.00E-35 | 46.15 | 255 | WP_042066964.1 |
| dihydrolipoyllysine-residue succinyltransferase [Escherichia coli] | Escherichia coli | 131 | 131 | 97% | 2.00E-35 | 46.15 | 252 | MDY9032137.1 |
| TPA: dihydrolipoyllysine-residue succinyltransferase [Escherichia coli] | Escherichia coli | 131 | 131 | 97% | 2.00E-35 | 46.15 | 252 | HBD1696035.1 |
| TPA: dihydrolipoyllysine-residue succinyltransferase [Escherichia coli] | Escherichia coli | 131 | 131 | 97% | 2.00E-35 | 46.15 | 250 | HBI3672065.1 |
| dihydrolipoyllysine-residue succinyltransferase [Escherichia coli] | Escherichia coli | 131 | 131 | 97% | 2.00E-35 | 46.15 | 253 | WP_347079825.1 |
| dihydrolipoyllysine-residue succinyltransferase [Escherichia coli] | Escherichia coli | 131 | 131 | 97% | 2.00E-35 | 46.15 | 253 | WP_136747018.1 |
| dihydrolipoyllysine-residue succinyltransferase [Escherichia coli] | Escherichia coli | 131 | 131 | 97% | 2.00E-35 | 46.15 | 253 | WP_136748022.1 |
| dihydrolipoyllysine-residue succinyltransferase [Escherichia coli] | Escherichia coli | 131 | 131 | 97% | 2.00E-35 | 46.15 | 253 | EFH6296858.1 |
| TPA: dihydrolipoyllysine-residue succinyltransferase [Escherichia coli] | Escherichia coli | 131 | 131 | 97% | 2.00E-35 | 46.15 | 254 | HDC0239447.1 |
| dihydrolipoamide succinyltransferase [Escherichia coli] | Escherichia coli | 131 | 131 | 97% | 2.00E-35 | 46.15 | 251 | NYZ48214.1 |
| dihydrolipoyllysine-residue succinyltransferase [Escherichia coli] | Escherichia coli | 131 | 131 | 97% | 2.00E-35 | 46.15 | 253 | WP_136801371.1 |
| dihydrolipoyllysine-residue succinyltransferase [Escherichia coli] | Escherichia coli | 131 | 131 | 97% | 2.00E-35 | 46.15 | 257 | MCU7719730.1 |
| dihydrolipoyllysine-residue succinyltransferase [Escherichia coli] | Escherichia coli | 131 | 131 | 97% | 2.00E-35 | 46.15 | 259 | EIC1734575.1 |
| TPA: dihydrolipoyllysine-residue succinyltransferase [Escherichia coli] | Escherichia coli | 131 | 131 | 97% | 2.00E-35 | 46.15 | 261 | HAM4331889.1 |
| dihydrolipoyllysine-residue succinyltransferase [Escherichia coli O177] | Escherichia coli O177 | 130 | 130 | 96% | 2.00E-35 | 46.11 | 229 | EFA8854754.1 |
| dihydrolipoyllysine-residue succinyltransferase [Escherichia coli] | Escherichia coli | 131 | 220 | 97% | 2.00E-35 | 46.15 | 260 | WP_252391012.1 |
| dihydrolipoyllysine-residue succinyltransferase [Escherichia coli] | Escherichia coli | 131 | 220 | 97% | 2.00E-35 | 46.15 | 259 | EGE6907964.1 |
| dihydrolipoyllysine-residue succinyltransferase [Escherichia coli] | Escherichia coli | 131 | 220 | 97% | 2.00E-35 | 46.15 | 259 | WP_119682542.1 |
| dihydrolipoyllysine-residue succinyltransferase [Escherichia coli] | Escherichia coli | 131 | 221 | 97% | 2.00E-35 | 46.15 | 263 | WP_047654718.1 |
| dihydrolipoamide succinyltransferase [Escherichia coli] | Escherichia coli | 131 | 221 | 97% | 2.00E-35 | 46.15 | 263 | OWF26202.1 |
| dihydrolipoamide succinyltransferase component (E2) [Escherichia coli] | Escherichia coli | 131 | 131 | 97% | 2.00E-35 | 46.15 | 269 | VFT70387.1 |
| hypothetical protein ECZU26_29390 [Escherichia coli] | Escherichia coli | 131 | 220 | 97% | 2.00E-35 | 46.15 | 260 | GHL32114.1 |

| | | | | | | | | |
|---|---|---|---|---|---|---|---|---|
| dihydrolipoyllysine-residue succinyltransferase [Escherichia coli] | Escherichia coli | 130 | 130 | 96% | 2.00E-35 | 46.11 | 230 | EGE7843444.1 |
| dihydrolipoyllysine-residue succinyltransferase [Escherichia coli O145:H28] | Escherichia coli O145:H28 | 131 | 220 | 97% | 2.00E-35 | 46.15 | 264 | EJH5270977.1 |
| TPA: dihydrolipoyllysine-residue succinyltransferase [Escherichia coli] | Escherichia coli | 131 | 220 | 97% | 2.00E-35 | 46.15 | 264 | HAN7662089.1 |
| TPA: dihydrolipoyllysine-residue succinyltransferase [Escherichia coli] | Escherichia coli | 131 | 220 | 97% | 2.00E-35 | 46.15 | 265 | HAP0635211.1 |
| dihydrolipoyllysine-residue succinyltransferase [Escherichia coli] | Escherichia coli | 130 | 130 | 95% | 3.00E-35 | 46.37 | 228 | WP_229024141.1 |
| 2-oxoglutarate dehydrogenase complex dihydrolipoyllysine-residue succinyltransferase [Escherichia coli] | Escherichia coli | 131 | 131 | 97% | 3.00E-35 | 46.15 | 269 | WP_115969540.1 |
| dihydrolipoamide succinyltransferase component (E2) [Escherichia coli] | Escherichia coli | 131 | 131 | 97% | 3.00E-35 | 46.15 | 269 | STG69599.1 |
| 2-oxoglutarate dehydrogenase complex dihydrolipoyllysine-residue succinyltransferase [Escherichia coli] | Escherichia coli | 131 | 131 | 97% | 3.00E-35 | 46.15 | 269 | WP_249524430.1 |
| 2-oxoglutarate dehydrogenase complex dihydrolipoyllysine-residue succinyltransferase [Escherichia coli] | Escherichia coli | 134 | 134 | 97% | 3.00E-35 | 46.7 | 405 | WP_097415199.1 |
| 2-oxoglutarate dehydrogenase complex dihydrolipoyllysine-residue succinyltransferase [Escherichia coli] | Escherichia coli | 131 | 131 | 97% | 3.00E-35 | 46.15 | 270 | WP_250382326.1 |
| 2-oxoglutarate dehydrogenase complex dihydrolipoyllysine-residue succinyltransferase [Escherichia coli] | Escherichia coli | 131 | 131 | 97% | 3.00E-35 | 46.15 | 280 | WP_320759799.1 |
| 2-oxoglutarate dehydrogenase complex dihydrolipoyllysine-residue succinyltransferase [Escherichia coli] | Escherichia coli | 134 | 224 | 97% | 3.00E-35 | 46.7 | 405 | MCW7185905.1 |
| 2-oxoglutarate dehydrogenase complex dihydrolipoyllysine-residue succinyltransferase [Escherichia coli] | Escherichia coli | 131 | 131 | 97% | 3.00E-35 | 46.15 | 275 | WP_136750337.1 |
| 2-oxoglutarate dehydrogenase complex dihydrolipoyllysine-residue succinyltransferase [Escherichia coli] | Escherichia coli | 131 | 131 | 97% | 3.00E-35 | 46.15 | 274 | MDY8964183.1 |
| 2-oxoglutarate dehydrogenase complex dihydrolipoyllysine-residue succinyltransferase [Escherichia coli] | Escherichia coli | 131 | 131 | 97% | 3.00E-35 | 46.15 | 274 | RIC38964.1 |
| 2-oxoglutarate dehydrogenase complex dihydrolipoyllysine-residue succinyltransferase [Escherichia coli] | Escherichia coli | 131 | 131 | 97% | 3.00E-35 | 46.15 | 279 | WP_247166683.1 |
| 2-oxoglutarate dehydrogenase complex dihydrolipoyllysine-residue succinyltransferase [Escherichia coli] | Escherichia coli | 131 | 131 | 97% | 3.00E-35 | 46.15 | 282 | MDY9040892.1 |
| 2-oxoglutarate dehydrogenase complex dihydrolipoyllysine-residue succinyltransferase [Escherichia coli] | Escherichia coli | 131 | 131 | 97% | 4.00E-35 | 46.15 | 286 | WP_047081876.1 |
| 2-oxoglutarate dehydrogenase complex dihydrolipoyllysine-residue succinyltransferase [Escherichia coli] | Escherichia coli | 131 | 131 | 97% | 4.00E-35 | 46.15 | 286 | WP_060643349.1 |
| 2-oxoglutarate dehydrogenase complex dihydrolipoyllysine-residue succinyltransferase [Escherichia coli] | Escherichia coli | 131 | 131 | 97% | 4.00E-35 | 46.15 | 295 | NAG00763.1 |

| | | | | | | | | |
|---|---|---|---|---|---|---|---|---|
| 2-oxoglutarate dehydrogenase complex dihydrolipoyllysine-residue succinyltransferase [Escherichia coli] | Escherichia coli | 131 | 131 | 97% | 4.00E-35 | 46.15 | 292 | WP_194498297.1 |
| 2-oxoglutarate dehydrogenase complex dihydrolipoyllysine-residue succinyltransferase [Escherichia coli] | Escherichia coli | 131 | 131 | 97% | 4.00E-35 | 46.15 | 289 | WP_250216047.1 |
| dihydrolipoyllysine-residue succinyltransferase [Escherichia coli] | Escherichia coli | 129 | 129 | 93% | 4.00E-35 | 46.29 | 226 | WP_074164196.1 |
| 2-oxoglutarate dehydrogenase complex dihydrolipoyllysine-residue succinyltransferase [Escherichia coli] | Escherichia coli | 131 | 131 | 97% | 4.00E-35 | 46.15 | 287 | EEZ2060122.1 |
| hypothetical protein EIMP300_69380 [Escherichia coli] | Escherichia coli | 128 | 128 | 93% | 5.00E-35 | 45.98 | 187 | BBU85538.1 |
| TPA: 2-oxoglutarate dehydrogenase complex dihydrolipoyllysine-residue succinyltransferase [Escherichia coli] | Escherichia coli | 132 | 132 | 97% | 5.00E-35 | 46.15 | 312 | HCO0761733.1 |
| TPA: 2-oxoglutarate dehydrogenase complex dihydrolipoyllysine-residue succinyltransferase [Escherichia coli] | Escherichia coli | 132 | 132 | 95% | 5.00E-35 | 46.07 | 318 | HCJ6193756.1 |
| 2-oxoglutarate dehydrogenase complex dihydrolipoyllysine-residue succinyltransferase [Escherichia coli] | Escherichia coli | 131 | 131 | 97% | 6.00E-35 | 46.15 | 303 | EEY6784672.1 |
| 2-oxoglutarate dehydrogenase complex dihydrolipoyllysine-residue succinyltransferase [Escherichia coli] | Escherichia coli | 131 | 131 | 97% | 6.00E-35 | 46.15 | 300 | MBA1840094.1 |
| 2-oxoglutarate dehydrogenase complex dihydrolipoyllysine-residue succinyltransferase [Escherichia coli] | Escherichia coli | 131 | 131 | 97% | 6.00E-35 | 46.15 | 315 | WP_284108106.1 |
| 2-oxoglutarate dehydrogenase complex dihydrolipoyllysine-residue succinyltransferase [Escherichia coli] | Escherichia coli | 132 | 132 | 95% | 6.00E-35 | 46.37 | 335 | MCF2049886.1 |
| 2-oxoglutarate dehydrogenase complex dihydrolipoyllysine-residue succinyltransferase [Escherichia coli] | Escherichia coli | 131 | 131 | 95% | 6.00E-35 | 46.07 | 318 | MBW2856955.1 |
| 2-oxoglutarate dehydrogenase complex dihydrolipoyllysine-residue succinyltransferase [Escherichia coli] | Escherichia coli | 133 | 133 | 97% | 6.00E-35 | 46.15 | 405 | WP_192459758.1 |
| TPA: 2-oxoglutarate dehydrogenase complex dihydrolipoyllysine-residue succinyltransferase [Escherichia coli] | Escherichia coli | 132 | 132 | 95% | 6.00E-35 | 46.37 | 336 | HDI5831311.1 |
| 2-oxoglutarate dehydrogenase complex dihydrolipoyllysine-residue succinyltransferase [Escherichia coli] | Escherichia coli | 131 | 221 | 97% | 7.00E-35 | 46.15 | 322 | WP_194154148.1 |
| 2-oxoglutarate dehydrogenase complex dihydrolipoyllysine-residue succinyltransferase [Escherichia coli] | Escherichia coli | 131 | 221 | 97% | 7.00E-35 | 46.15 | 322 | MCU8601182.1 |
| 2-oxoglutarate dehydrogenase complex dihydrolipoyllysine-residue succinyltransferase [Escherichia coli] | Escherichia coli | 131 | 221 | 97% | 7.00E-35 | 46.15 | 321 | EEY6693670.1 |
| TPA: 2-oxoglutarate dehydrogenase complex dihydrolipoyllysine-residue succinyltransferase [Escherichia coli] | Escherichia coli | 132 | 132 | 95% | 7.00E-35 | 46.37 | 327 | HDH9429852.1 |
| 2-oxoglutarate dehydrogenase complex dihydrolipoyllysine-residue succinyltransferase [Escherichia coli] | Escherichia coli | 131 | 131 | 97% | 7.00E-35 | 46.15 | 324 | EER8331471.1 |
| 2-oxoglutarate dehydrogenase complex dihydrolipoyllysine-residue succinyltransferase [Escherichia coli] | Escherichia coli | 131 | 131 | 97% | 7.00E-35 | 46.15 | 324 | MDO4358214.1 |

| | | | | | | | | |
|---|---|---|---|---|---|---|---|---|
| 2-oxoglutarate dehydrogenase complex dihydrolipoyllysine-residue succinyltransferase [Escherichia coli] | Escherichia coli | 132 | 222 | 97% | 7.00E-35 | 46.15 | 331 | WP_042068848.1 |
| TPA: 2-oxoglutarate dehydrogenase complex dihydrolipoyllysine-residue succinyltransferase [Escherichia coli] | Escherichia coli | 131 | 131 | 95% | 8.00E-35 | 46.37 | 328 | HDH2834759.1 |
| dihydrolipoamide succinyltransferase component (E2) [Escherichia coli] | Escherichia coli | 132 | 222 | 97% | 8.00E-35 | 46.15 | 332 | STM59242.1 |
| 2-oxoglutarate dehydrogenase complex dihydrolipoyllysine-residue succinyltransferase [Escherichia coli] | Escherichia coli | 133 | 133 | 97% | 8.00E-35 | 46.7 | 405 | EEZ9068234.1 |
| 2-oxoglutarate dehydrogenase complex dihydrolipoyllysine-residue succinyltransferase [Escherichia coli] | Escherichia coli | 133 | 223 | 97% | 8.00E-35 | 45.6 | 405 | ELT8823651.1 |
| 2-oxoglutarate dehydrogenase complex dihydrolipoyllysine-residue succinyltransferase [Escherichia coli] | Escherichia coli | 131 | 221 | 97% | 8.00E-35 | 46.15 | 318 | MCX0506393.1 |
| 2-oxoglutarate dehydrogenase complex dihydrolipoyllysine-residue succinyltransferase [Escherichia coli] | Escherichia coli | 133 | 133 | 97% | 9.00E-35 | 46.15 | 405 | EKM5464437.1 |
| 2-oxoglutarate dehydrogenase complex dihydrolipoyllysine-residue succinyltransferase [Escherichia coli] | Escherichia coli | 131 | 221 | 97% | 9.00E-35 | 46.15 | 328 | WP_229002757.1 |
| TPA: 2-oxoglutarate dehydrogenase complex dihydrolipoyllysine-residue succinyltransferase [Escherichia coli] | Escherichia coli | 131 | 221 | 97% | 9.00E-35 | 46.15 | 332 | HCN8464961.1 |
| 2-oxoglutarate dehydrogenase complex dihydrolipoyllysine-residue succinyltransferase [Escherichia coli] | Escherichia coli | 131 | 221 | 97% | 9.00E-35 | 46.15 | 337 | WP_048224949.1 |
| 2-oxoglutarate dehydrogenase complex dihydrolipoyllysine-residue succinyltransferase [Escherichia coli] | Escherichia coli | 131 | 221 | 97% | 1.00E-34 | 46.15 | 338 | MDY8712251.1 |
| 2-oxoglutarate dehydrogenase complex dihydrolipoyllysine-residue succinyltransferase [Escherichia coli] | Escherichia coli | 133 | 133 | 97% | 1.00E-34 | 46.7 | 405 | EFL9867405.1 |
| 2-oxoglutarate dehydrogenase complex dihydrolipoyllysine-residue succinyltransferase [Escherichia coli] | Escherichia coli | 131 | 131 | 97% | 1.00E-34 | 46.15 | 345 | MBA1060735.1 |
